# Pattern and component responses of primate MT neurons recorded with multi-contact electrodes, stimulated with 1D and 2D noise patterns

**DOI:** 10.1101/2021.12.23.474004

**Authors:** Christian Quaia, Incheol Kang, Bruce G Cumming

## Abstract

Direction selective neurons in primary visual cortex (area V1) are affected by the aperture problem, i.e., they are only sensitive to motion orthogonal to their preferred orientation. A solution to this problem first emerges in the middle temporal (MT) area, where a subset of neurons (called pattern cells) combine motion information across multiple orientations and directions, becoming sensitive to pattern motion direction. These cells are expected to play a prominent role in subsequent neural processing, but they are intermixed with cells that behave like V1 cells (component cells), and others that do not clearly fall in either group. The picture is further complicated by the finding that cells that behave like pattern cells with one type of pattern, might behave like component cells for another.

We recorded from macaque MT neurons using multi-contact electrodes while presenting both type I and unikinetic plaids, in which the components were 1D noise patterns. We found that the indices that have been used in the past to classify neurons as pattern or component cells work poorly when the properties of the stimulus are not optimized for the cell being recorded, as is always the case with multi-contact arrays. We thus propose alternative measures, which considerably ameliorate the problem, and allow us to gain insights in the signals carried by individual MT neurons.

We conclude that arranging cells along a component-to-pattern continuum is an oversimplification, and that the signals carried by individual cells only make sense when embodied in larger populations.

## Introduction

In primates, unlike in rodents, neither the retina nor the lateral geniculate nucleus contain direction-selective cells. Such cells are first found in the primary visual cortex (area V1), where they represent a minority of the cells (around 20%, De Valois, Yund, and Hepler, 1982; Hawken, Parker, and Lund, 1988; Hamilton, Albrecht, and Geisler, 1989; Prince et al., 2002). It is in the middle temporal (MT) area, which is the recipient of direct (Cragg, 1969; Maunsell and van Essen, 1983a) and fast-conducting (Ungerleider and Mishkin, 1979; Van Essen, Maunsell, and Bixby, 1981) projections from area V1, that motion signals first become ubiquitous (Allman and Kaas, 1971; Dubner and Zeki, 1971; Maunsell and Van Essen, 1983b; Albright, 1984). Based on this observation, and on the effects of causal manipulations of the activity of its neurons (Salzman et al., 1992), area MT is thus widely considered crucial in processing visual motion information. Furthermore, thanks to observations from many electrophysiological experiments, we now have a reasonable understanding of which signals are carried by MT neurons, and how they are generated from their V1 inputs, at least for simple visual stimuli (Rust et al., 2006).

Neurons in area V1 are affected by the so-called *aperture problem*: a spatio-temporal filter that is narrowband for orientation lets through only motion energy in a direction orthogonal to that orientation. Accordingly, V1 direction-selective cells respond only to the component of image motion orthogonal to their preferred orientation. In MT, many neurons also behave like this, and are called *component* cells. However, one observation that indicates that area MT is specialized for motion processing is that some of its neurons appear to solve the aperture problem: MT *pattern* cells integrate information across different orientations to correctly signal the motion direction of broadband patterns (Movshon et al., 1985; Movshon and Newsome, 1996). This specialization is usually revealed using stimuli composed of two sinusoidal gratings (called *plaids*), which are narrowband in spatial frequency. However, models describing pattern cells postulate an integration of signals over both spatial frequency and orientation (Simoncelli and Heeger, 1998; Perrone, 2004; Rust et al., 2006; Nishimoto and Gallant, 2011; Quaia, Optican, and Cumming, 2016).

Here we report neurophysiological data from macaque MT cells stimulated with plaids obtained by summing two one-dimensional noise patterns, which we call random line plaids, previously used in behavioral experiments in humans (Quaia, Optican, and Cumming, 2016; Quaia et al., 2019). The use of these plaids offers a significant advantage when recording from multiple neurons simultaneously, as there is no need to adjust the spatial frequency content of the stimulus to optimally activate individual neurons. However, since it is not possible to generate a stimulus that matches the different preferred speeds of the neurons being recorded, the stimulus is inevitably too slow for some cells and too fast for others. As we will show, this severely limits the usefulness of the pattern index (Movshon et al., 1985; Smith, Majaj, and Movshon, 2005; Rust et al., 2006), the measure most widely used to distinguish component and pattern cells. We will further show that the cross-correlation measure recently proposed to characterize MT cells based on their response to unikinetic plaids (Wallisch and Movshon, 2019) does not always work well either.

After demonstrating the shortcomings of current measures, we will propose some alternatives, which capture the extent to which responses of individual MT cells reflect a pattern motion computation even when suboptimal stimuli are presented.

## Materials and Methods

### Electrophysiology

We recorded extracellular spiking activity from two male rhesus monkeys (*macaca mulatta*). Surgery under general anesthesia was performed in each monkey to implant a head post and a recording chamber over a craniotomy in correspondence of area MT on one side of the brain. During each recording session the monkey was awake and had its head restrained by means of a head post. While the monkey passively fixated a central cross (for which it was periodically rewarded with drops of water or fruit juice), stimuli were presented in rapid succession and neural activity was recorded from area MT using multi-contact linear electrodes (V-probes, Plexon, 24 contacts with a spacing of 50 *μm*). Analog electrical signals from each electrode were digitized at 40 *kHz* and stored to disk for off-line analysis.

All procedures were performed in accordance with the U.S. Public Health Service Policy on the humane care and use of laboratory animals. All protocols were approved by the National Eye Institute Animal Care and Use Committee.

### Spike sorting

Only well isolated units were used for the analyses reported here. This required careful spike sorting of the signals recorded from the multi-contact electrodes. Since fully automated spike sorting methods do not currently have the required accuracy, we used a custom-developed interactive method, in which an initial automated process was manually verified and refined by an experienced user. Ambiguous cases were discussed by two users and if a consensus was not reached the putative unit in question was discarded.

### Visual stimuli

The data presented here were part of a larger study, under which several types of visual stimuli were presented while recording from area MT. Here we focus on the responses evoked in MT neurons by four classes of stimuli:

- A one-dimensional (1-D) random line pattern, having one of 6 orientations, and drifting in one of 12 directions (always orthogonal to its orientation), spaced uniformly around the circle (i.e., 30° apart). Stimulus direction is assigned according to the following convention: In a 0° 1-D noise pattern the lines are vertical and drift to the right. Angles increase in a counter-clockwise direction (90° indicates upward motion, 180° leftward and 270/ – 90° downward). The drifting speed orthogonal to the orientation of the lines was 10 in some sessions and 15°/*s* in the others. Note that these stimuli are narrowband in orientation and speed, but broadband in spatial (SF) and temporal frequency (TF).
- A random dot stimulus (RDS, 2-D noise), drifting at the same speed and in the same directions as the 1-D noise patterns above. These stimuli are broadband in orientation, SF, and TF, and are only narrowband in speed.
- Plaids obtained by summing two 1-D noise patterns, having directions of motion 120° apart and drifting at the same speed. The second 1-D pattern is rotated 120° clockwise relative to the first one. We define as direction of motion for this class of stimuli the direction of motion of the first of the two components. Thus, in a 0° type I plaid the first component is a 0° 1-D noise pattern (i.e., it has vertical lines drifting to the right) and the second is a 240/ – 120° 1-D noise pattern (i.e., the lines are rotated by 60° counter-clockwise relative to the first component, and drift down and to the left). Note that the motion direction of the stimulus as a rigid object (the so-called pattern motion direction) is –60° (down and to the right, half-way between the direction of motion of the two components). In Figure 1 we graphically represent these stimuli using two solid arrows, pointing in the directions of motion of the two drifting 1-D noise patterns.
- Unikinetic plaids, obtained by combining a drifting 1-D noise pattern (same speed and directions as the 1-D noise patterns presented in isolation described above) and a static 1-D noise pattern that is rotated 45° counter-clockwise (we indicate this configuration as UP45) or clockwise (UP-45) relative to the drifting 1-D noise pattern, which defines the direction of motion of the stimulus. For example, in a 0° UP45 plaid, the first component is a 0° drifting 1-D noise pattern (i.e., vertical lines drifting to the right) and the second is a 45° static 1-D noise pattern (i.e., its lines slant from top left to bottom right). The pattern motion direction of the plaid as a whole is parallel to the orientation of the static 1-D component, with the direction determined by the drifting component. For a 0° UP45 plaid the pattern motion direction is thus –45°, down and to the right. In Figure 1 we graphically represent these stimuli using a solid arrow, pointing in the direction of motion of the drifting 1-D component, and a solid line, parallel to the orientation of the static 1-D component.

**Figure 1.**
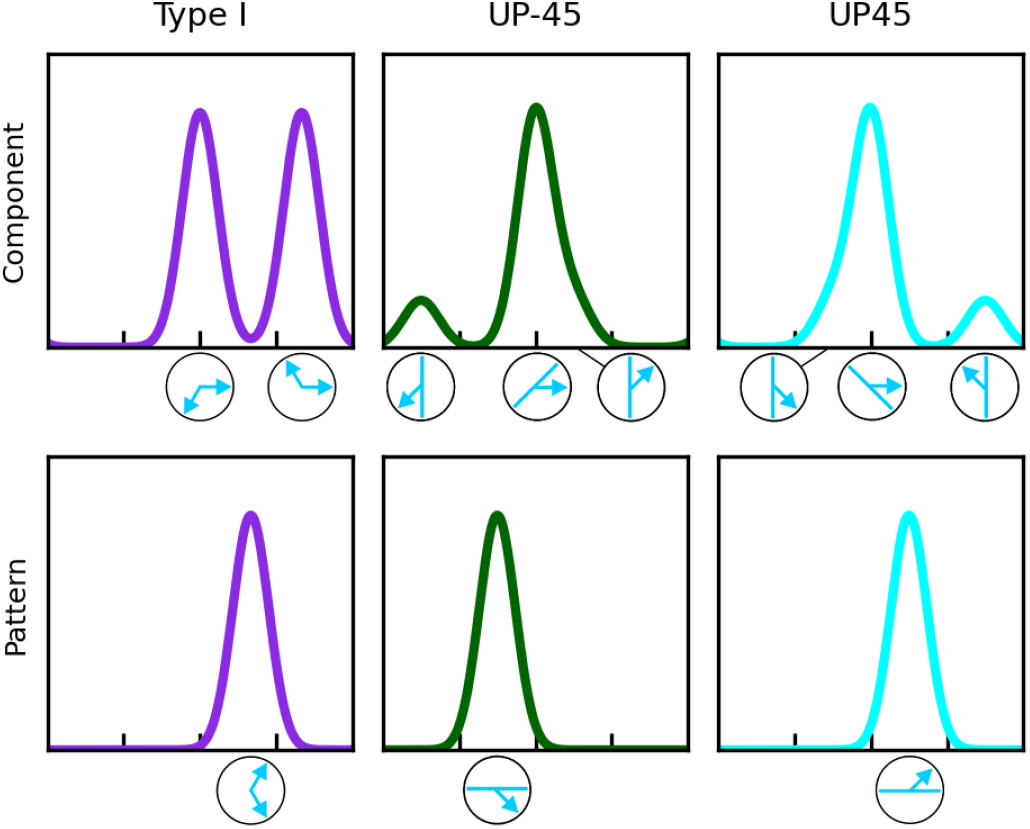
Response of idealized component and pattern cells, preferring motion to the right, to the plaid stimuli we presented. **Top row**: The response of an ideal component cell to a plaid is equal to the sum of its responses to each individual component (an output nonlinearity could be expected to follow the sum, but it is not usually considered). The 1-D components of each stimulus at key stimulus directions are indicated graphically under the abscissa. The responses to unikinetic plaids must consider the response of the cell to the both drifting (arrows) and static (solid lines) components. Their relative strength might vary considerably from cell to cell. **Bottom row**: An ideal pattern cell will respond to the direction of motion of the plaid as if it were a rigid object. It should thus respond maximally when the plaid as a whole appears to drift to the right.

The stimuli were randomly intermixed and presented to the animal in rapid succession (500 *ms* presentations separated by 100 *ms* intervals during which the screen was blank at mean luminance, 27.9 *cd/m*^2^). The stimulus was always presented in a circular aperture, as if drifting behind it. The size and location of the aperture were fixed in a recording session, and varied from session to session as a function of the location and size of the receptive fields (RF) of the units being recorded. Unlike classical single-unit recordings, the size of the stimuli used was thus not matched to the classical RF size of each neuron, but almost invariably extended into their RF surround. An even more important difference between our study and previous studies in MT is that whereas in single-unit studies the stimulus speed (as well as other parameters) is at least approximately matched to the preferred speed of the neuron, with array recordings this is not possible. For all neurons recorded in any one session we thus used the same drifting speed, selected to be in the center of the previously reported distribution of preferred speeds for MT neurons (Wang and Movshon, 2016). By recording responses to both 1-D and 2-D noise stimuli we could however infer, at least for some units, when our stimulus was too slow or too fast relative to the preferred speed of a neuron being recorded. Theoretical considerations (Kawakami and Okamoto, 1996; Simoncelli and Heeger, 1998) and previous studies (Okamoto et al., 1999; Kumano and Uka, 2013) indicate that the direction tuning curve for a 1-D stimulus that is too slow for a MT pattern cell will be bimodal; similarly, the direction tuning curve for a 2-D noise stimulus that is too fast for a MT component cell will also be bimodal. As expected, in our recordings we identified several such bimodal tuning curves to 1-D or 2-D noise stimuli.

### Data fitting and interpolation

We quantified the tuning properties of our neuronal sample by computing for each neuron the single-trial spike count in a 500 *ms* time window, starting 50 *ms* after stimulus onset and ending 50ms after stimulus offset. This time window was selected because the response latency of individual cells ranged between 35 and 50 *ms*. We recorded from each neuron during multiple (mean: 18.7; median: 16) presentations of each stimulus. Mean tuning curves were then computed by averaging the spike count over all presentations of the same stimulus. The 68% confidence interval for each mean spike count was computed using standard bootstrap-based non-parametric techniques. The resting firing rate of each neuron was estimated by counting all the spikes generated in the period from 50 *ms* before to 30 *ms* after the onset of a stimulus (a period which should exclude most stimulus-evoked activity).

Direction tuning curves have traditionally been fitted to data using a Von Mises (i.e., a circular Gaussian) function. This function would however not be appropriate for many of our tuning curves, which as noted above are often not unimodal. Various fitting or interpolating functions (polynomial, spline, Lowess, etc.) could then be used, but we reasoned that, given the circular periodicity of the tuning curves, the simplest fitting function is a smoothed version of a Fourier interpolator. Because we took 12 samples around the circle, the Fourier transform has seven harmonics (mean response, *H*_1_, *H*_2_, *H*_2_, *H_3_, *H**_4_, *H*_6_, and *H_6_*, the last corresponding to the Nyquist frequency). To smooth it, we simply set to zero the amplitude of the two highest frequency harmonics (i.e., *H*_5_ and *H_6_*, those with a period of 72° and 60°, respectively). Note that, unlike other fitting schemes, this one is deterministic, and has no free parameters. We verified its ability to also fit well unimodal responses by using a Von Mises function to generate responses with various fullwidth at half-height (FWHH) bandwidths (between 20° and 180°), sampling them every 30° (matching the spacing in our experiments) starting with a random phase, and comparing the mean-squared error of our Fourier fit to that of a Von Mises fit. We found that our Fourier method performs as well or better than the Von Mises fit whenever the FWHH is larger than 40° (the Von Mises function should in principle always be able to fit perfectly, given that the testing function is derived from it, but the fitting procedure does not always converge to the global minimum given the sparse sampling), but it cannot generate narrower curves (remember that we smoothed it by removing the two highest SF components). With narrow tuning curves (FWHH less than 30°) the Von Mises fit was often very accurate in terms of its error, but the parameters of the function could be quite different from those of the generating function (because with such curves only two samples are sizable, the fitting procedure often converges to a very narrow and tall Von Mises function, whose peak is much larger than any of the data points, a problem that could be partially addressed through regularization of the fitting function). Even for unimodal tuning curves, a Von Mises fit is thus rarely more reliable or appropriate than a smoothed Fourier interpolation. Accordingly, for all figures we used the Fourier interpolator to display the data, and used Von Mises fits only in the context of tuning curve bandwidth estimation).

The FWHH bandwidth of unimodal tuning curves was estimated by measuring the span between the angles at which the fitted tuning curve dropped from its peak to half its peak, on both sides of the peak. For bimodal tuning curves, when the trough between the two peaks was smaller than the half-height, we took the span of the outer half-height angles as measures of the FWHH bandwidth. We started off by fitting all functions with our smoothed Fourier interpolator; for tuning curves in which the computed FWHH was less than 35°, we then refit the tuning curve using a regularized Von Mises function, and used this function to estimate the FWHH.

We estimated the preferred direction of each cell from the vector average of the tuning curve associated with drifting 1-D or 2-D noise stimuli^2^. Visual inspection of the tuning curves revealed that the 2-D noise tuning curve usually provides a more reliable estimate of the preferred direction of MT neurons, and we thus by default used these stimuli. For the (component) cells in which the direction tuning curve to 2-D noise stimuli was bimodal (see above), we however used the 1-D noise direction tuning curve instead. While theoretically the two peaks of the bimodal 2-D noise direction tuning curve should straddle the preferred direction, and thus the vector average should still provide a reliable measure, we found that in a significant subset of cases the two peaks were quite asymmetric, and the 1-D noise preferred direction provided a more accurate estimate (as determined from the responses to the various plaids). We also used the direction estimate based on 1-D noise rather than 2-D noise whenever the response to the best 1-D noise stimulus was more than three times as large as that to the best 2-D noise stimulus, a rare occurrence in our dataset.

### Spike-count correlations

To compute the signal correlation between two units, we first computed, for each unit, the average spike count for each stimulus (only trials during which both units were well isolated were considered), and then computed the Pearson’s correlation between the two spike counts for the two cells. To compute the noise correlation between two units we first z-scored the single-trial spike counts separately for each stimulus and cell, and then computed the overall Pearson’s correlation between the z-scores (across all stimuli) for the two cells. Fisher’s r-to-Z transformation was used to compute population means and standard deviations of the correlation values.

## Results

In previous studies, two indices have been used to functionally classify MT cells into component and pattern types:

- The pattern index (*PI*), which is based on the cell’s responses to type I plaids and to their 1-D components;
- The unikinetic rotation (*UR*), which is based on the cell’s responses to two sets unikinetic plaids, having the same moving component but differently oriented static components.

As we now show, both indices are useful, but they have limits, especially when applied to our data. To be able to highlight their merits and limits, and to develop and validate new measures, it is useful to identify *a priori* component and pattern cells. Accordingly, one of us (C.Q.), based on visual inspection of their tuning curves to all stimuli, divided our 386 cells into five groups:

- *Component-fast*: 63 cells that behaved mostly as expected from component cells, but had a bimodal tuning curve to 2-D noise, indicating that the stimulus was too fast relative to their preferred speed;
- *Component*: 101 cells that behaved mostly as a component cell would be expected to;
- *Mixed*: 70 cells which represented neither pattern nor component velocity, or behaved like pattern cells for one type of plaids but as component cells for another;
- *Pattern*: 56 cells that behaved mostly as a pattern cell would be expected to;
- *Pattern-slow*: 96 cells that behaved mostly as pattern cells, but their direction tuning curve to 1-D noise was bimodal, indicating that the stimulus was too slow relative to their preferred speed.

This exercise revealed that, while some cells are clearly more pattern-like or component-like (although cells that perfectly match the ideal models are rare), for many the attribution is unclear, and making one’s criterion looser or stricter could significantly change the fraction of cells classified as “mixed”. Nevertheless, cells that were classified as pattern certainly did not behave as component, and vice-versa. Thus, any index or measure used to classify the cells should not mix up cells in those two groups.

### Pattern Index (*PI*)

The pattern index compares the direction tuning curve measured with type I plaids to those predicted for ideal component and pattern cells (Figure 1), based on the responses to the components of the plaid. An ideal component cell is assumed to respond to the two 1-D stimuli (whether sinusoidal gratings or noise patterns) present in a plaid independently. Its direction tuning curve to a type I plaid is thus the sum of its 1-D component direction tuning curve and the same direction tuning curve rotated by 120° in a counter-clockwise (i.e., positive) direction. An ideal pattern cell is instead assumed to respond to the pattern direction of the plaid stimulus as if it were a rigid object; its tuning curve for a type I plaid is thus equal to the 1-D component direction tuning curve, rotated by 60° (i.e., halfway between the two plaid components) in a counter-clockwise direction.

The pattern index (Rust et al., 2006) computes the relative similarity of the shape (magnitude information is discarded) of the measured type I plaid direction tuning curve to these two model tuning curves. For each model, the partial correlation between measured and model responses is computed, and Fisher’s r-to-Z transform is applied, converting the value to a Z-correlation. The pattern index is then defined as the difference between the Z-correlation for the pattern and that for the component model. Negative values indicate that responses to type I plaids are more similar to the component prediction, whereas positive values indicate a closer match to the pattern prediction. Values between −1.28 and 1.28 (corresponding to a *p* = 0.9 significance level) are difficult to interpret, and can indicate that neither prediction is accurate, or that both are equally accurate (either because the cell response falls in between those predicted by the two models, or because the two predictions are not very different, which happens when the 1-D component tuning curve is broad). No class is assigned to cells whose pattern index falls in this range. In previous studies this occurred in about a third of the cells. As we’ll now show, in our dataset the situation is more complicated.

In Figure 2A we plot the Z-correlation for the component and pattern models (with models for the type I plaid tuning curves based on the response to the drifting 1-D noise patterns) for our cells, color-coded according to the five groups that were manually identified. It is apparent that most component cells (blue dots) are in the lower right corner, and thus are correctly classified by the pattern index. Similarly, most gray dots, indicating cells that were hard to manually identify as either pattern or component cells, fall around the main diagonal in the plot, and are thus unclassed by the pattern index. However, most of the orange dots, corresponding to the pattern-slow group, are not in the upper left corner, where pattern cells belong. In Figure 2B we make this clear by plotting the distribution of the pattern index for the five groups. The pattern index correctly classifies most component cells; it does a reasonable job even with mixed cells (most are unclassed). However, it incorrectly classifies many pattern cells as either component or unclassified, and it does an especially poor job with the pattern-slow group. To quantify its performance, we trained a binary linear classifier on the pattern index and applied it to the 316 cells manually identified as either component (fast or normal) or pattern (slow or normal). The pattern index correctly classified 74.3% (performance from stratified 10-fold cross-validation) of the samples (threshold set at −1.48), with *d*′ = 1.11.

**Figure 2.**
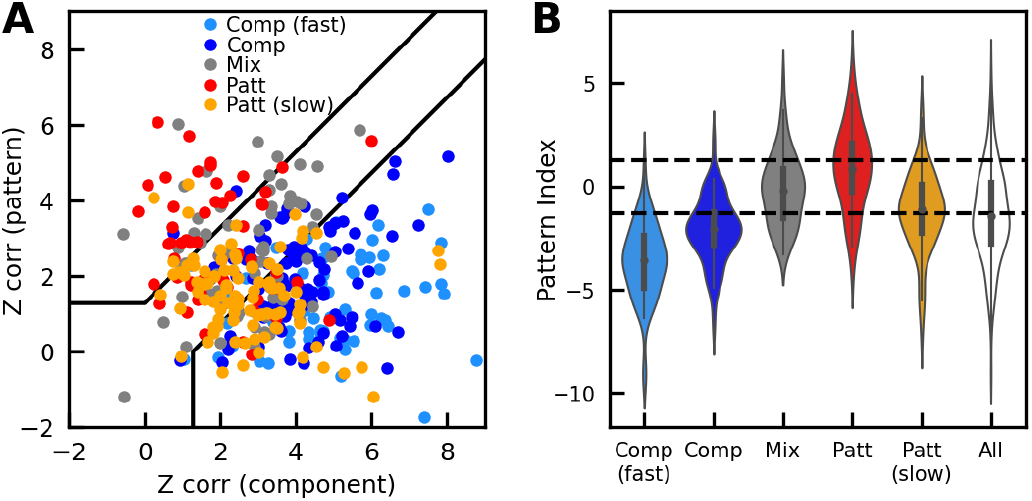
Classification performance of the pattern index (*PI*). **A**: For each cell recorded we indicate the partial Z-correlation with component (abscissa) and pattern (ordinate) models. Each cell is color-coded according to its manually assigned membership in one of five groups (legend). **B**: Distribution of the pattern index (difference between the partial Z-correlation for pattern and component models) for each group of cells. The dashed lines indicate the −1.28 and 1.28 significance levels.

This relatively low performance (chance is at 50%) can be attributed to two factors. First, when pattern cells respond to a 1-D stimulus that is too slow relative to their preferred speed (an unavoidable occurrence with array recordings), their tuning curve is bimodal (with the two peaks usually straddling the actual preferred direction of the cell, as estimated from the response to 2-D noise stimuli). In most of these cells, the type I plaid tuning curve is however unimodal, or it has a large primary peak and smaller secondary peaks, and the ideal pattern cell prediction (i.e., a rotated bimodal curve) is thus a poor match.

In Figure 3 we show tuning curves for five representative cells from this group (the tuning curves have been rotated so that each cell’s preferred direction, measured with drifting 2-D noise, is zero). It is obvious that, for all five cells, the strongest response to a type I plaid is observed around 60°, and that the direction tuning curve for 1-D noise patterns is bimodal, two signatures of a pattern cell for which the 1-D stimulus is moving too slowly. As we will discuss later, the responses to unikinetic plaids (last row) also point to these as being pattern cells. On the other hand, it is also obvious that the type I plaid tuning curve (purple) is a poor match for the pattern model (red dashed lines) based on the drifting 1-D noise direction tuning curve; in fact, it better matches the component model (blue dashed line), especially for the first three cells. It is thus no wonder that, according to the pattern index, these units would be (incorrectly) classified as component cells. Thus, for these cells the pattern index, as computed, is misleading.

**Figure 3.**
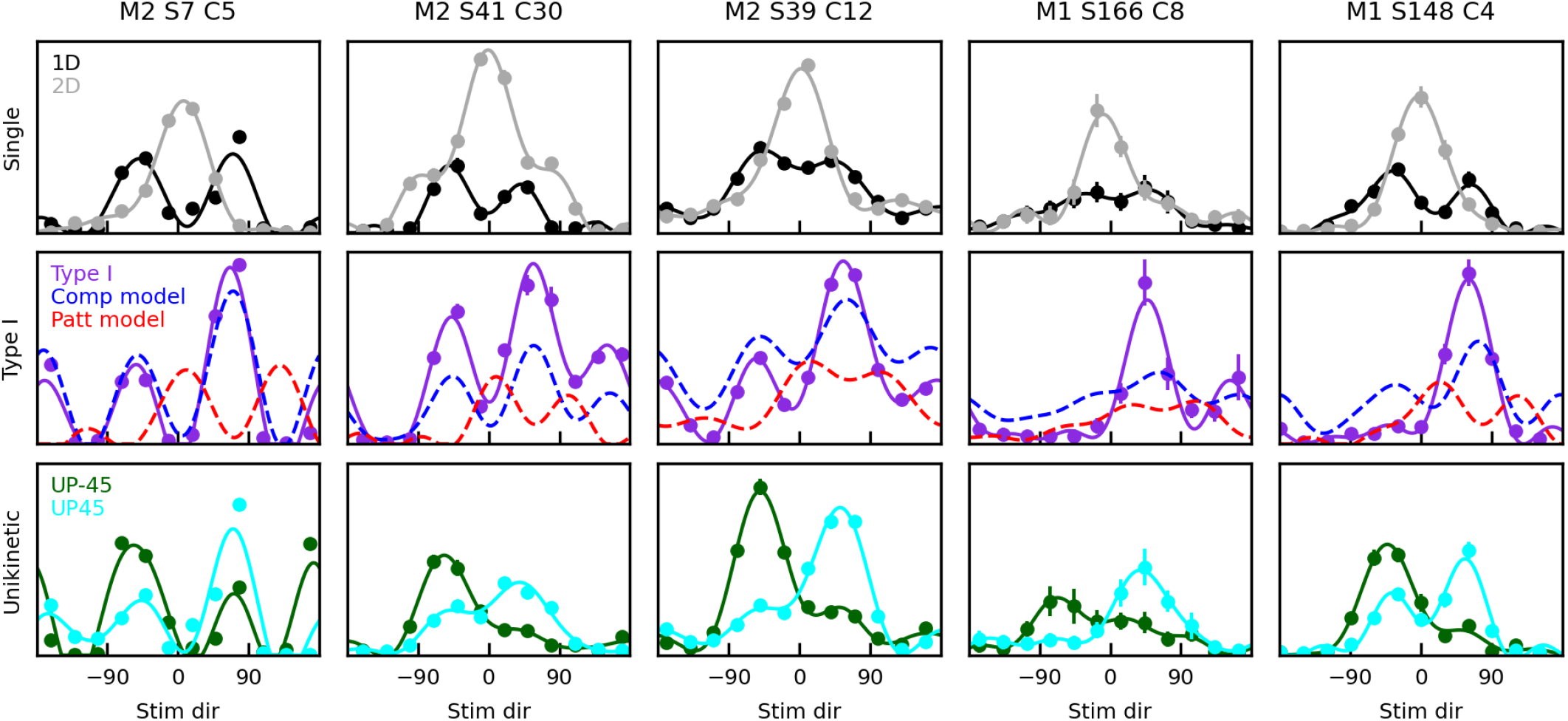
Tuning curves for five cells from the pattern-slow group. Responses to 1-D (black) and 2-D (gray) drifting noise patterns are shown in the top row. Responses to type I noise plaids (purple) are shown in the middle row, together with the predictions (based on the response to 1-D noise patterns) of pattern (dashed red) and component (dashed blue) models. Responses to unikinetic plaids (UP-45 in green, UP45 in cyan) are shown in the bottom row. For actual responses, the average and its 68% confidence interval is indicated for each data point, and the solid curves are Fourier-based interpolants (see Methods). For model responses, only Fourier-based interpolants are shown. Values of all pattern indices described in the paper for the cells shown (here and in later figures) are listed in the Appendix.

The second problem with the pattern index arises because in some cells the preferred direction for 1D noise is quite different from that for 2D stimuli (three examples are shown in Figure 4)^3^. In these cases, using the response to 1D noise as the basis for the component/pattern class assignment is misleading - examination of the responses invariably reveals that the preferred direction based on the 2-D noise stimuli is the correct one. As in Figure 3, the strongest response to a type I plaid is observed 60° from the preferred direction of the cell, as identified by the 2-D noise direction tuning curve (and as expected from a pattern cell). And yet, the pattern index is negative, because the component prediction based on 1-D noise better fits the type I plaid tuning curve than the pattern prediction.

**Figure 4.**
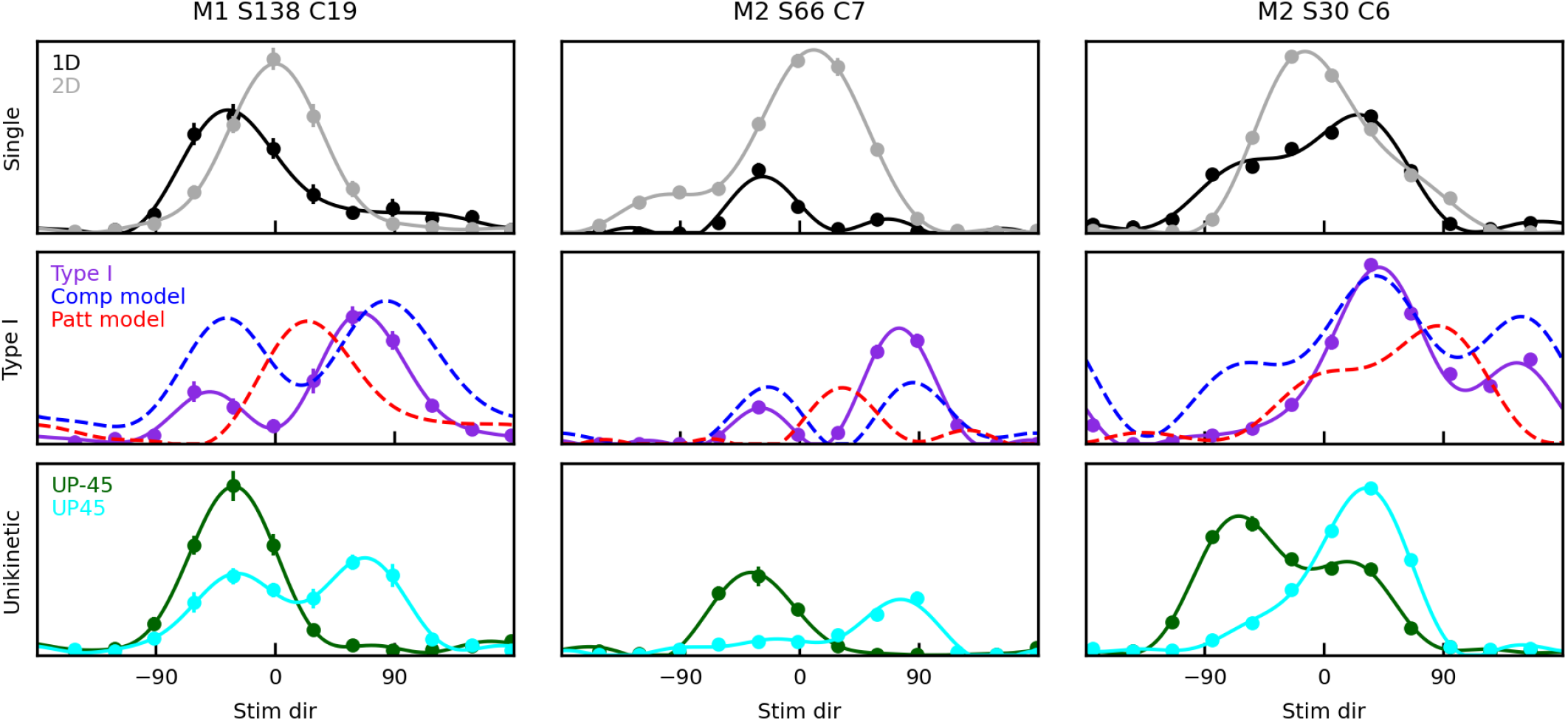
Tuning curves for three sample cells in which the preferred direction for 1-D noise and 2-D noise stimuli was significantly different. Same conventions as in Figure 3.

### Unikinetic Rotation (*UR*)

The unikinetic rotation (*UR*) index (Wallisch and Movshon, 2019) estimates the relative rotation (in degrees) between the tuning curves of two families of unikinetic plaids (in our study UP45 and UP-45). It is computed by locating the peak of the cross-correlation function of the two tuning curves. With our convention, component cells should have a *UR* close to zero, whereas pattern cells should have a UR close to 90°. Because of the limited and variable response of MT neurons to static stimuli (Albright, 1984; Wallisch and Movshon, 2019), this measure is more reliable than an index, analogous to the pattern index, that compares the responses to the plaid and its components used in a previous study (Khawaja, Liu, and Pack, 2013) and illustrated in Figure 1 (bottom row).

With directions sampled every 30°, simply looking for the maximum of the cross-correlation between the two direction tuning curves would results in an unacceptably coarse (30°) resolution. Accordingly, the *UR* is typically computed as a weighted average of the cross-correlation function values (i.e., a center-of-mass/vector average operation, preferably on r-to-Z transformed values).

In Figure 5A we show the distribution of the *UR* index for our cells, color-coded according to the five groups that were manually identified. In Figure 5B we show the classification of cells, based on the UR (there is no established way of doing this; we chose to classify as component cells those with *UR* < 30°, as pattern cells those with *UR* > 60°, and as unclassed those with intermediate *UR* values) and on our manual criterion. As desired, the unikinetic rotation classifies most component cells as either component or unclassed, most pattern cells as pattern or unclassed, and most mixed cells as unclassed. However, almost half of the cells end up being unclassed; furthermore, some of the rotations are very large (either positive or negative) and hard to interpret. To quantify the performance of the *UR* index, we again trained a binary linear classifier on the unikinetic rotation and applied it to the 316 cells manually identified as either component (fast or normal) or pattern (slow or normal). The unikinetic rotation correctly classified 75.6% (performance from stratified 10-fold cross-validation) of the samples (threshold set at 42.49°), with *d*′ = 1.16. It thus performs similarly to the pattern index.

**Figure 5.**
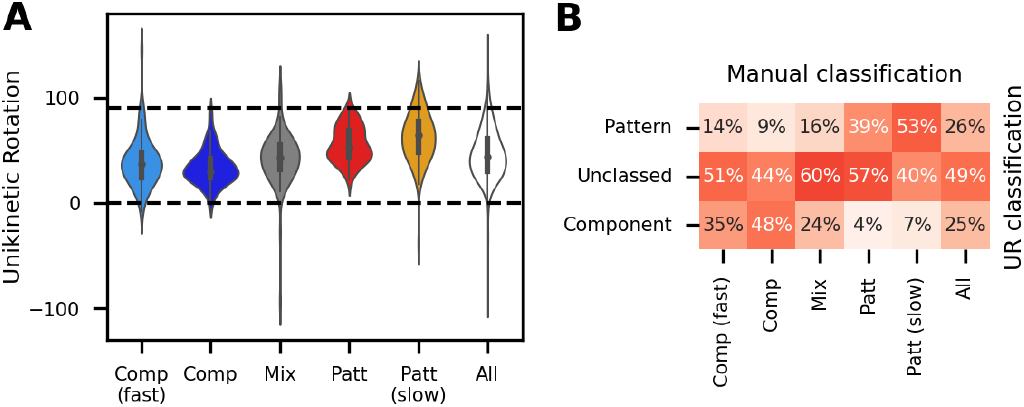
Classification performance of the unikinetic rotation index (*UR*). **A**: Distribution of the unikinetic rotation index for each group of cells. The dashed lines indicate zero (rotation associated with an ideal component cell) and 90° (rotation associated with an ideal pattern cell). **B**: Classification matrix. Different columns are associated with different groups of neurons, based on our manual classification; rows indicate instead the classification based on unikinetic rotation (Component if *UR* < 30°, Pattern if *UR* > 60°. Unclassed otherwise). The percentage of cells in each intersection of manual and *UR*-based groups is indicated and color-coded (darker means higher).

One possibility is that in fact with unikinetic plaids the rotation is only partial, and that the measure is thus correctly capturing the behavior of the cells. What we’d then expect is that the direction tuning curves for the two unikinetic plaids would be in between those shown in Figure 1 for ideal component and pattern cells. They would thus have a single large peak, whose location would be offset by more than 0° but less than 90°. However, this is rarely observed in cells that have been manually classified as either pattern or component. Instead, the direction tuning curves often have two peaks, and the major peak in one usually coincides with the secondary peak of the other (e.g., Figures 3 and 4, bottom row). Our manual classification was mostly based on the relative shift of the major peak.

In pattern cells, this often results in a cross-correlation function that has its major peak at large positive rotations, but from there drops off faster for larger than for smaller rotations. For pattern cells, an estimate of unikinetic rotation based on a center-of-mass interpolation is then biased towards zero relative to the peak. Some examples are shown in Figure 6, together with the interpolated cross correlation function (bottom row). In more extreme cases there is an actual peak of the cross-correlation function at zero rotation, and the unikinetic rotation is then close to 45° (Figure 7). This is obviously misleading, since the tuning curves to the two unikinetic plaids (third row) are far from being simply a rotated version of each other.

**Figure 6.**
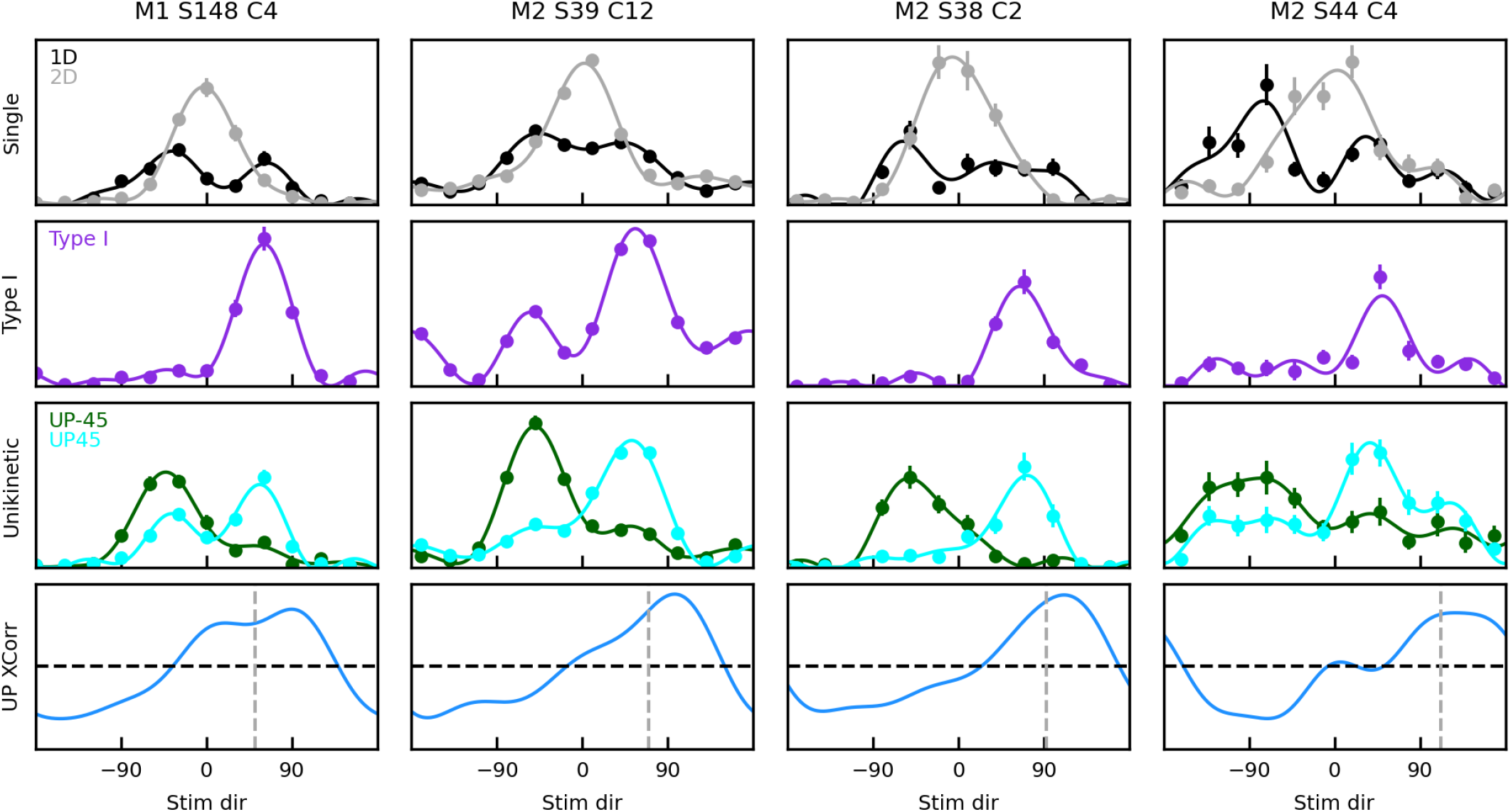
Tuning curves for four cells (manually classified as pattern-slow cells) in which the cross-correlation between the UP45 and UP-45 tuning curves (bottom row) peaks around 90°, but is skewed towards zero, resulting in a reduced unikinetic rotation estimate. The dashed horizontal line indicates the zero cross-correlation level; the dashed gray line indicates the center-of-mass cross-correlation value (i.e., *UR*). Direction tuning curves (average and its 68% confidence interval for each data point, and Fourier-based interpolant) for 1-D and 2-D noise patterns (top row), type I plaids (second row), unikinetic plaids (UP45 and UP-45, third row) are also shown.

**Figure 7.**
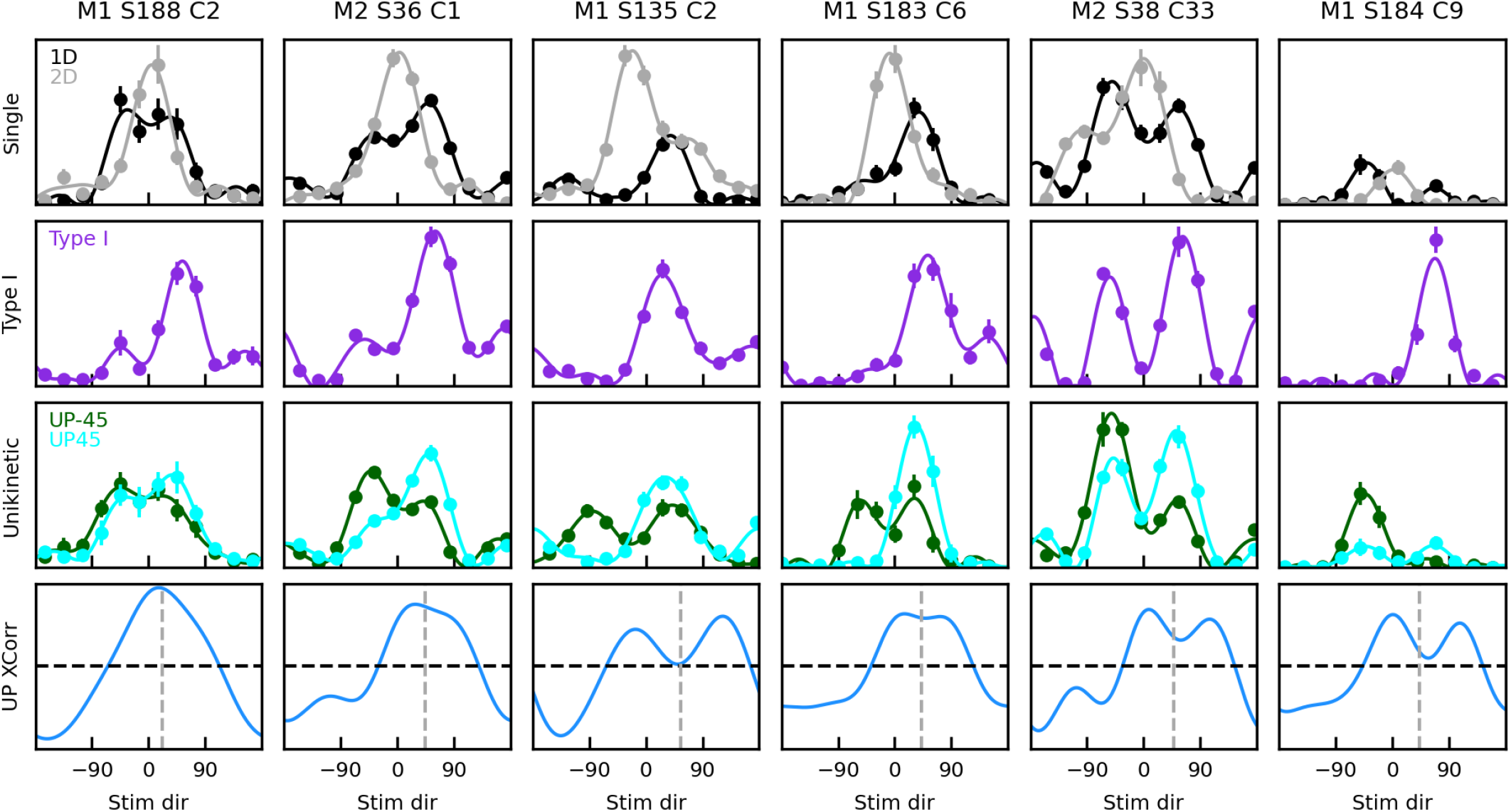
Tuning curves for six cells (manually classified as pattern-slow cells) in which the cross-correlation between the UP45 and UP-45 tuning curves (bottom row) has a peak around 0°, resulting in a unikinetic rotation estimate (gray dashed line) biased towards zero. Same convention as in Figure 6.

Note that all cells in Figures 6 and 7 have UP45 direction tuning curves with clear peaks at positive angles, and UP-45 direction tuning curves with clear peaks at negative angles, and that the two peaks are separated by at least 90°. However, both tuning curves also have a secondary peak at the opposite sign rotation. What differentiates the cells in Figure 6 from those in Figure 7 is that in the former group the peak at the ”wrong” direction is, for both tuning curves, considerably smaller than that at the ”correct” rotation. In contrast, for the cells in Figure 7 in at least one of the two tuning curves that two peaks are of comparable magnitude, resulting in a peak around zero rotation in the cross-correlation function. But what is the origin of these peaks? As noted, these cells belong to the pattern-slow group, and accordingly their direction tuning curve to a drifting 1-D noise pattern is bimodal. Inspection of the figures reveals that the two peaks of this direction tuning curve (top row, black) coincide with the peaks in the tuning curves of the unikinetic plaids (third row). The presence of the static component in the unikinetic plaids thus results in an enhancement or suppression of each of these two peaks, the extent of which ends up determining the unikinetic plaid tuning curves, and thus their cross-correlation function. In an ideal pattern cell, the positive (i.e., counter-clockwise) peak should be enhanced in the UP45 tuning curve and suppressed in the UP-45 tuning curve; the opposite should happen to the negative (i.e., clockwise) peak. To understand to what extent this holds, in Figure 8 we plot, for cells belonging to the pattern-slow group, the pairwise distribution of five measures: the base-2 logarithm of the ratio between or UP-45) and the 1-D noise pattern at two orientations the (interpolated) response to an unikinetic plaid (UP45 (identified, for each cells, as the orientation between 0° and 100° where the UP45 tuning curve peaks, and the orientation between −100° and 0° where the UP-45 tuning curve peaks), and the unikinetic rotation. We labeled the measures as follows:

- **P45**: log_2_(UP45[*pos_pk_*]/1D[*pos_pk_*])
- **N45**: log_2_(UP45[*neg_pk_*]/1D[*neg_pk_*])
- **P-45**: log_2_(UP-45[*pos_pk_*]/1D[*pos_pk_*])
- **N-45**: log2(UP-45[*neg_pk_*]/1D[*neg_pk_*])

**Figure 8.**
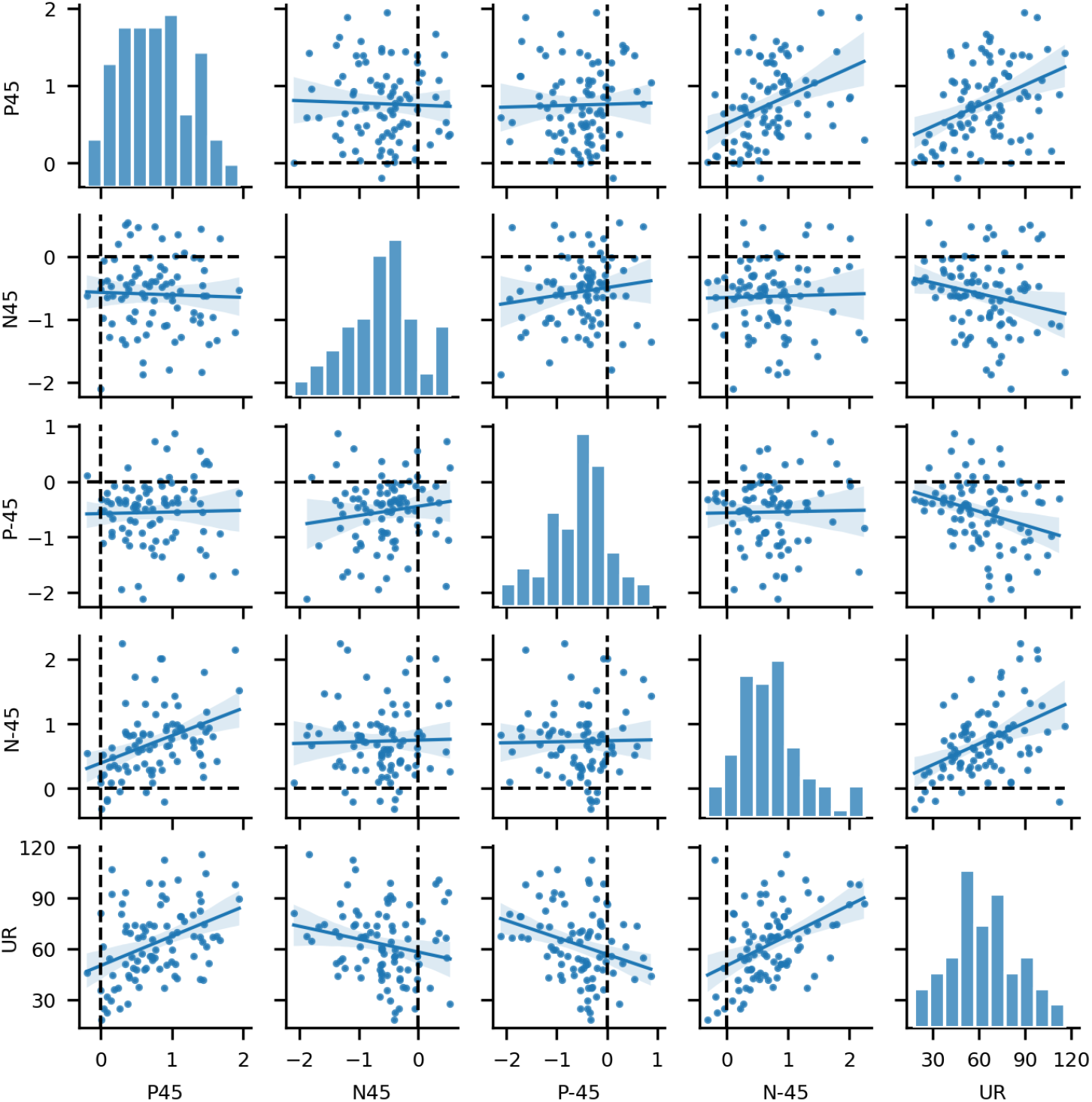
Paired relationship between estimates of enhancement/suppression for unikinetic plaids in the directions corresponding to the peaks in the 1-D noise tuning curve, for cells manually classified as pattern-slow. Their relation to the unikinetic rotation index is also shown (last row/column). In pattern cells, P45 and N-45 (first and third rows and columns) should both be positive, whereas P-45 and N45 (second and third rows and columns) should be negative. This is true for most cells. All four measures are also significantly correlated (*p* < 0.05) with the UR (last row/column).

To compensate for the unreliability of the ratio when dealing with small values, we dropped data-points that were more than 2 standard deviations away from the mean (7 data points out of 96).

This analysis confirms the inferences we had made above from the examples shown in Figures 6 and 7. First of all, note that all four measures have the expected sign for most cells, indicating that both the expected suppression and the expected enhancement are present in the population. The two measures of enhancement (P45 and N-45) are positively and significantly correlated with each other; the two measures of suppression (N45 and P-45) are positively, but not significantly, correlated. Finally, the measures are all significantly correlated with the unikinetic rotation (with the expected sign). Thus, the problem with many pattern-slow cells is that there is usually a clear peak in the cross-correlation function at 90°, but it is not dominant enough, resulting in an unikinetic rotation measure that is considerably biased towards smaller rotations.

As we will now show, in component cells, and especially so in the component-fast group, we have the opposite problem: The major peak of the cross-correlation function at or near zero is almost invariably accompanied by a secondary peak around 135°, which induces a positive bias in the unikinetic rotation measure. The origin of this secondary peak can be inferred from Figure 1 (top row): It is the result of matching the primary peak in one tuning curve, with the side peak due to the static pattern in the other (located around 135° for UP45 plaids, and around −135° for UP-45 plaids, see Figure 1, top row). Some examples are shown in Figure 9. Sometimes the secondary peaks in the direction tuning curves are large enough that what should be the secondary peak of the cross-correlation function is in fact the primary, and the peak at zero becomes secondary (Figure 10).

**Figure 9.**
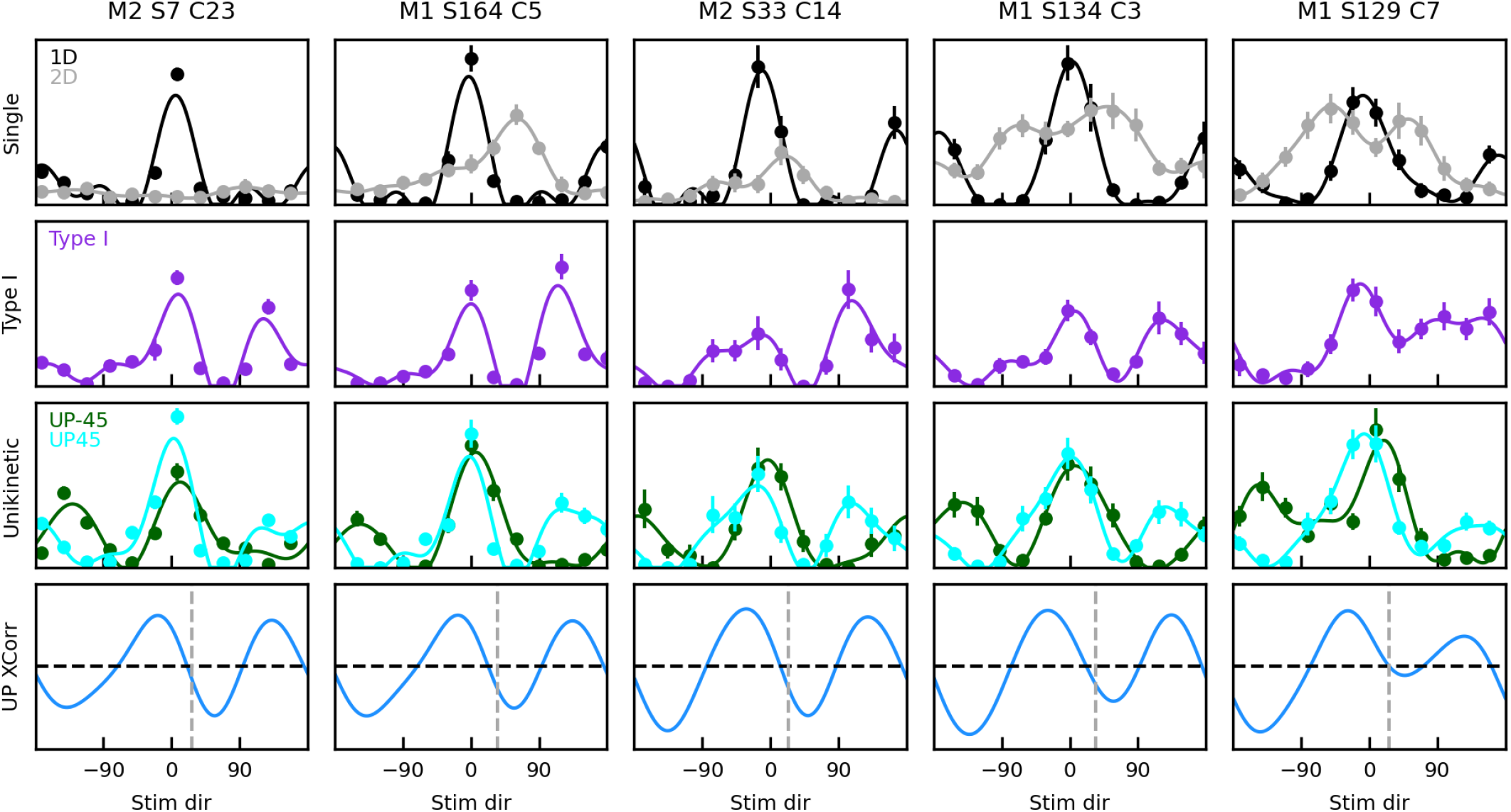
Tuning curves for five cells (manually classified as component or component-fast) in which the cross-correlation between the UP45 and UP-45 tuning curves (bottom row) has a major peak around 0°, but a strong secondary peak around 120° – 135°, resulting in a positively biased unikinetic rotation estimate (gray dashed line). Same convention as in Figure 6.

**Figure 10.**
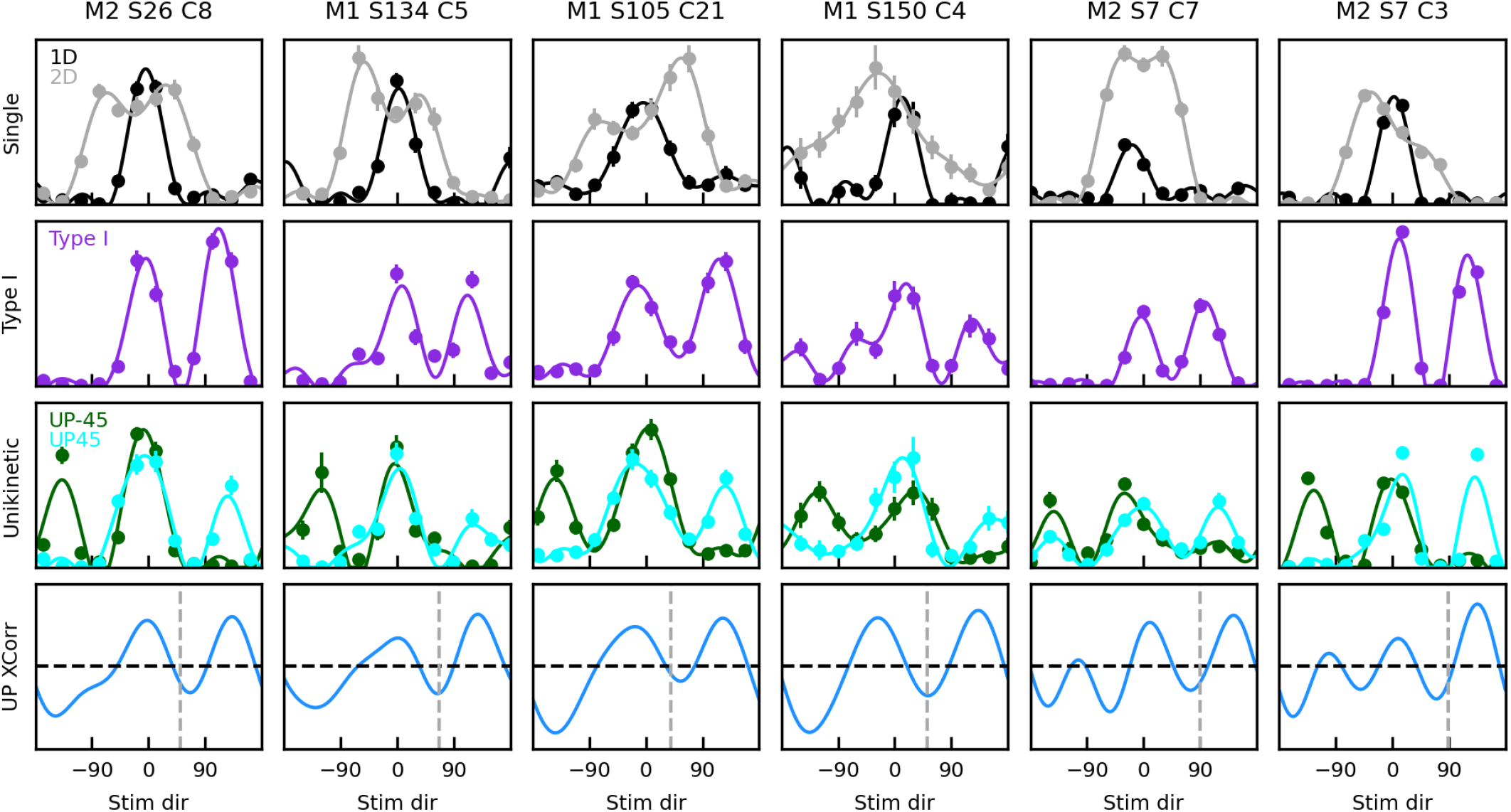
Tuning curves for six cells (manually classified as component or component-fast) in which the cross-correlation between the UP45 and UP-45 tuning curves (bottom row) has a peak around 0°, but an even larger peak around 120° – 135°, resulting in a strongly positively biased unikinetic rotation estimate. Same convention as in Figure 6.

When stimuli are optimized for the preferred speed of each cell, the pattern index and the unikinetic rotation are positively correlated (Wallisch and Movshon, 2019). However, in our dataset they are not (*r* = 0.044, *p* = 0.391, dark green line Figure 11). Combining them only slightly improves the classification performance: The cross-validated classification performance of a linear support vector classifier trained on the component and pattern cells is 79.7% (the solid black line in Figure 11 shows the optimal linear class boundary), with *d*′ = 1.8.

**Figure 11.**
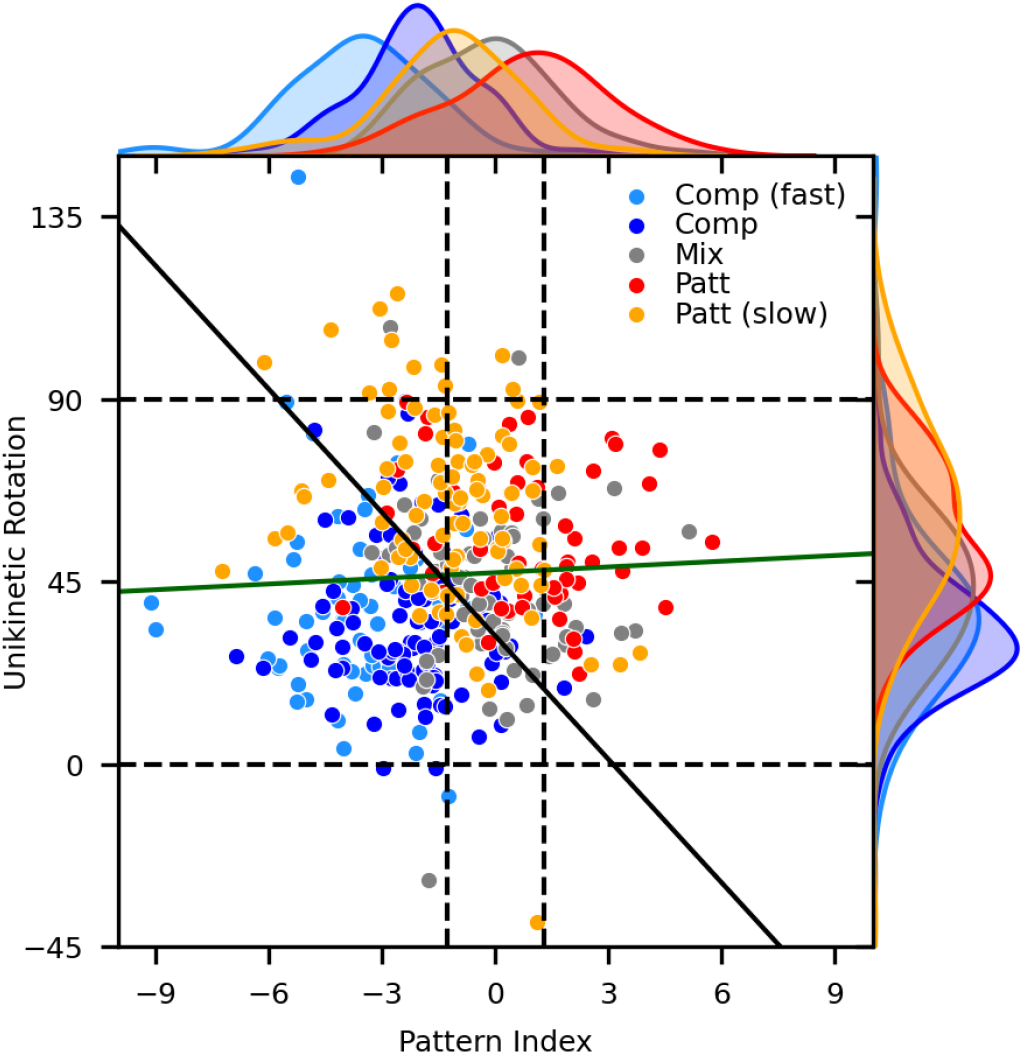
The pattern index and the unikinetic rotation both carry information about the nature of each cell (see Figure 2 and 5), but they are not correlated across all cells (dark green line, *r* = 0.044, *p* = 0.391). Marginal distributions (top and right) indicate distinct distributions for the five groups, and thus an undesired sensitivity to suboptimal stimulus speed. Their individual performance can be slightly improved by combining them into a single index, for example by training a binary linear Support Vector Classifier (excluding the Mixed group). The solid black line indicates the optimal boundary of this classifier (cells to the right are classified as pattern, to the left as component).

### Population responses

The examples we have shown above indicate that the direction tuning curves observed at the single-cell level are very diverse. If we consider subpopulations of cells it is however conceivable that part of this variability will cancel out. In Figure 12 we plot the tuning curves for all the cells, grouped according to their manual group assignment, and rotated to align their preferred direction with zero. Each tuning curve was also independently z-scored. Importantly, to provide a modicum of averaging over similar cells, the plot was smoothed by using a boxcar filter across sequences of 20 units. All the issues that we have discussed above are still present in this population view, but the smoothing has considerably ameliorated them. This of course has also the side-effect of smoothing the transitions between groups.

**Figure 12.**
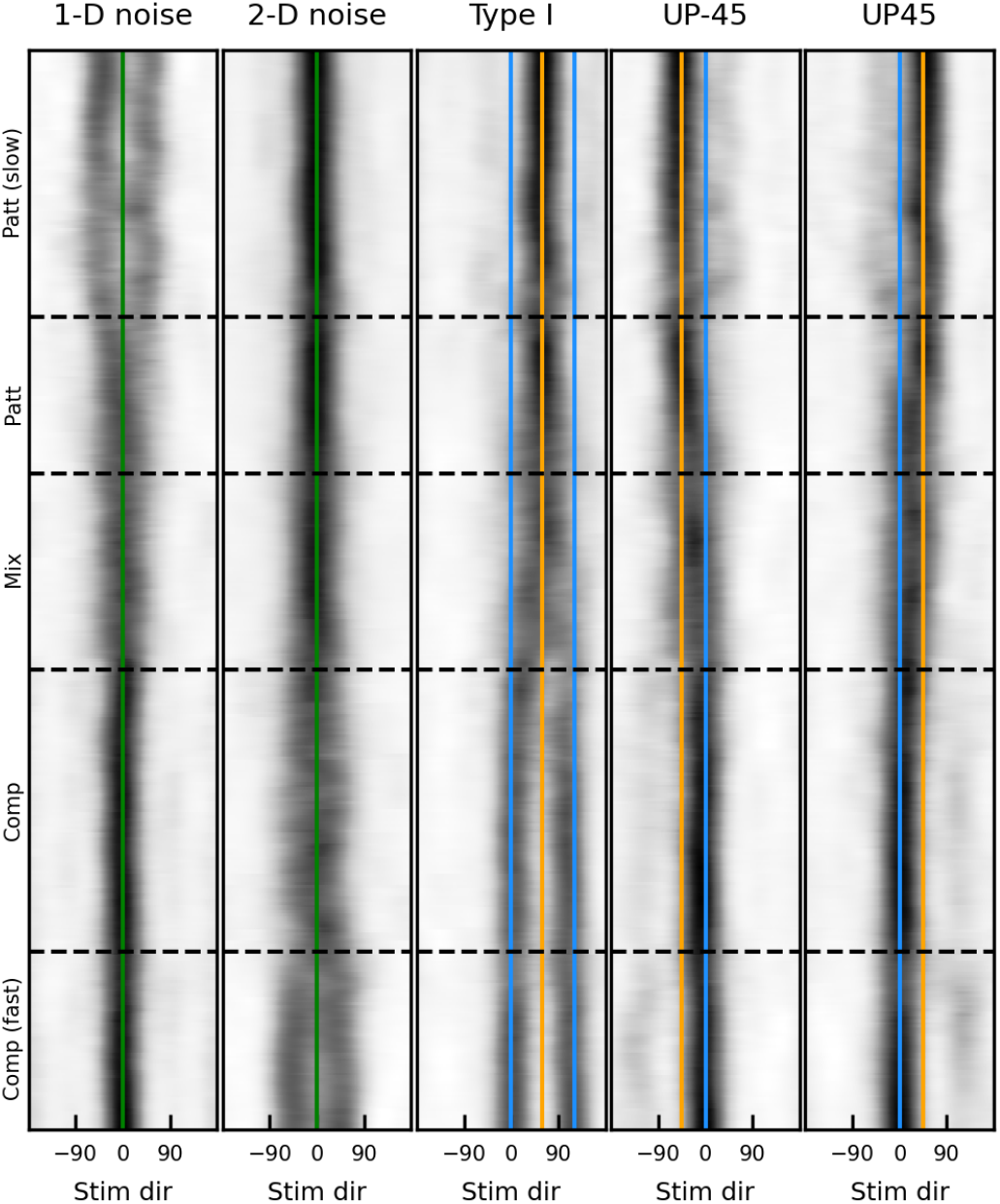
Direction tuning curves for all our cells for each stimulus type, after aligning their preferred direction with 0° and z-scoring them. The cells are grouped according to our manual classification, and within each group they are sorted according to our newly proposed *gPI* index. A boxcar filter is used to smooth local variability over 20 cells. Green lines: 0° direction; blue lines: predicted peak location for component cells; orange lines: predicted peak location for pattern cells.

We see that the pattern-slow group has bimodal tuning curves to the 1-D noise stimulus, but a unimodal response, centered at 60° for type I plaids, with only a hint of the responses at other directions that we observed in Figure 3.

Even for cells in the pattern group we see that there is more directional scatter in 1-D noise responses compared to those to 2-D stimuli. These are the issues that impaired the performance of the pattern index. This population view makes clear that in pattern-slow cells the direction tuning curve to type I plaids resembles a rotated (by 60° counter-clockwise) version of the direction tuning curve to 2-D, not 1-D, noise patterns. The problems that plagued the unikinetic rotation are also still visible, although attenuated by the smoothing. For example, we see that in component and component-fast cells the tuning curves to unikinetic plaids peak at 0°, but there are also some side peaks at ±135°, as we saw in some sample cells shown in Figures 9 and 10. Conversely, in pattern cells the major peak for unikinetic plaids is around ±45°, but there are secondary peaks in the ∓45° directions (as we saw in Figure 7). There are indeed some cells that have a bona-fide unikinetic intermediate rotation, but those belong to the mixed group, and possibly spill over in the pattern group (to which they were probably assigned because of their clear peak response at 60° for type I plaids).

To better visualize the directional signal that is carried by these subpopulations, in Figure 13 we show, for each stimulus type, the mean (and 68% confidence interval) of these population tuning curves, separately for each group.

**Figure 13.**
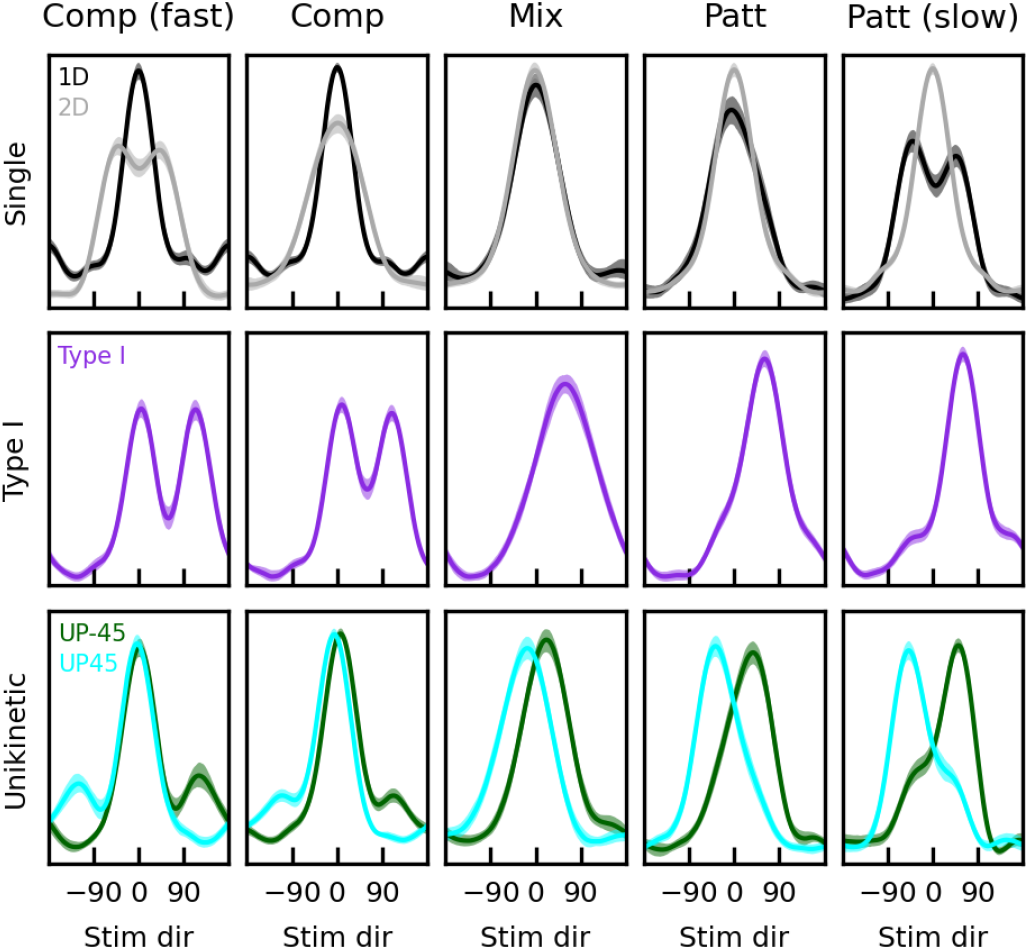
Average of the preferred-direction aligned and z-scored direction tuning curves for all stimuli, separately for each manual classification group. The 95% confidence interval of each tuning curve is plotted (shading).

### Developing alternative indices

The above analysis demonstrates that current indices are not well suited to characterize MT cells when they are recorded using frequency broadband stimuli drifting at suboptimal speeds. With simultaneous recordings becoming standard, this problem will only become more common. One approach to address this issue would be to modify the indices, for example by first identifying cells in the pattern-slow and component-fast groups (based on the bimodality of direction tuning curves to 1-D or 2-D stimuli, respectively) and treating them differently. The alternative is to define new indices, ideally based on those aspects of the responses of component and pattern cells that are robust to variations induced by stimulus speed. For ease of usage, it would also be desirable to have bounded indices, for example yielding a value close to −1 for an ideal component cell and close to +1 for an ideal pattern cell (this is not the case with the indices discussed in the previous sections).

Given the periodic nature of the tuning curves, we reasoned that their first five Fourier harmonics *H*_0_ ⋯ *H*_4_ could be good candidates to build new indices. As can be seen in the many examples shown above, our smoothed Fourier interpolant captures the direction tuning curves very well. It has 10 parameters (which we indicate as *A*_0_ ⋯ *A*_4_ and *P*_0_ ⋯ *P*_4_, where *A* stands for amplitude and *P* for phase of the corresponding Fourier coefficient *F_i_*) for each tuning curve, which can be reduced to 8 by dropping *A*_0_ (the mean of the tuning curve, which correlates with the peak firing rate of the cell, and it’s irrelevant for our purposes), and *P*_0_ (always zero). Furthermore, harmonics with a small amplitude are bound to be sensitive to noise, and thus unlikely to be useful. In our dataset, for all tuning curves the amplitude generally decreases as frequency increases. For unikinetic plaids, the means of *A*_0_ ⋯ *A*_4_ are [80, 75, 42, 26, 14]; with type I plaids they are [100, 88, 33, 40, 18]. If we exclude the fourth harmonic, we are thus left with just three harmonics, 6 parameters in all.

In Figure 14 we plot, for each of the within-group mean z-scored direction tuning curves shown in Figure 13, their tuning curves (Full) and the first three harmonics of their Fourier decomposition (*H*_1_, *H*_2_, and *H*_3_), for each of the plaid stimuli used.

**Figure 14.**
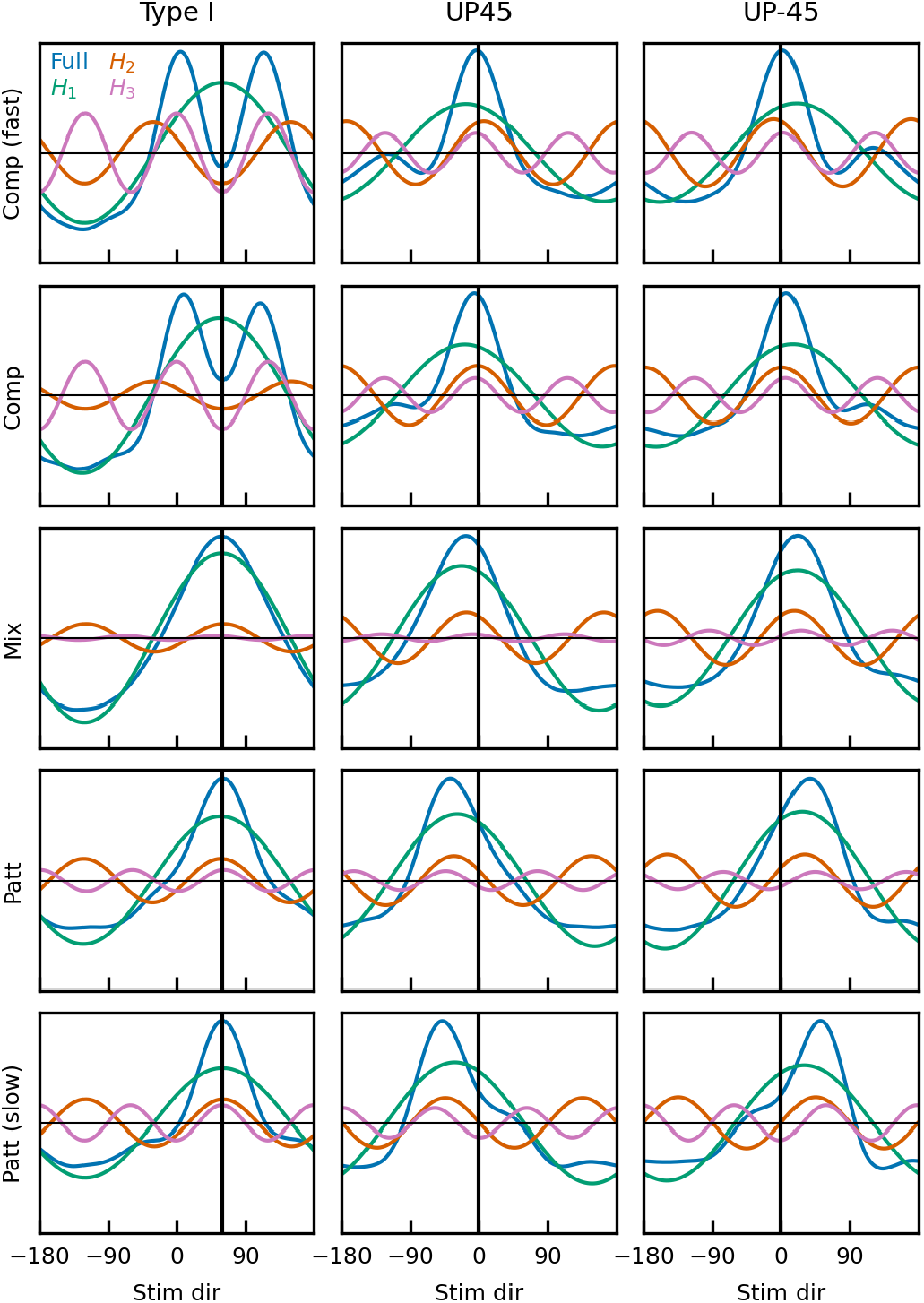
Relationship between the top three Fourier harmonics for the group-specific averages shown in Figure 13. The relationship between the phase of *H*_1_ (green) and that of *H*_2_ (red) and *H*_3_ (pink) varies across groups for type I plaids, and can thus be used for classification purposes. Similarly, the phase of *H*_2_ varies across groups for UP45 and UP-45 plaids.

### Bikinetic Pattern Index (*bPI*)

With type I plaids (Figure 14, left column), the phase of *H*_1_ (green lines) is the same in all cell groups, peaking at 60°. In pattern cells, *H*_1_, *H*_2_ (red lines), and *H*_3_ (pink lines) peak together, and a single peak at 60° emerges in the tuning curve. In component cells, *H*_2_ and *H*_3_ have a trough were *H*_1_ peaks, and as a result the tuning curve is bimodal. The difference in phase between *H*_1_ and *H*_2_ (or *H*_3_) can thus be used to discriminate component from pattern cells. Of course, these are only average cells, but they provide a clear indication that the phase of *H*_2_ and *H*_3_ where *H*_1_ peaks could carry information about pattern identity (essentially by discriminating a unimodal from a bimodal type I tuning curve). With elementary algebra we obtain that for *H*_2_ this phase is *P*_2_ – 2*P*_1_, and for *H*_3_ it is *P*_3_ – 3*P*_1_.

The phase values for the first three harmonics can be very simply obtained using the matrix form of the discrete Fourier transform. If we indicate with ***y*** the column vector with the average spike counts of a tuning curve, and define a row vector of integers ***x*** = [0 ⋯ 11], we have that the complex coefficients of the first three harmonics (neglecting scaling factors) are:

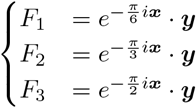

Amplitude and phase for these three complex coefficients can then be easily extracted (in Matlab and Python/Numpy with the *abs* and *angle* functions).

In Figure 15A, we plot, for all cells (color-coded according to our manual classification, as in Figure 11), these two phase differences. As expected from Figure 14, in pattern cells (red and orange) both phases have values around zero, whereas in component cells (blue and cyan) both have values around ±180° (we have constrained angles to the ±180° range). We can thus define an index based on each of these two measures:

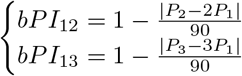

*bPI* stands for bikinetic (since there are two drifting 1D patterns in a type I plaid) pattern index, and the subscripts identify the harmonics used. Each index varies between −1 (putative component cell) and 1 (putative pattern cell). In Figure 15B we plot these two indices. They are strongly correlated (*r* = 0.728, *p* = 5.8 × 10^−65^), but provide some amount of complementary information. We could thus combine them in a single index. Because the sensitivity to noise of a harmonic phase will be proportional to its amplitude, we simply compute a weighted sum of these two measures based on the amplitude of the respective harmonics:

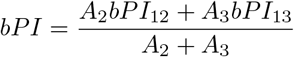

**Figure 15.**
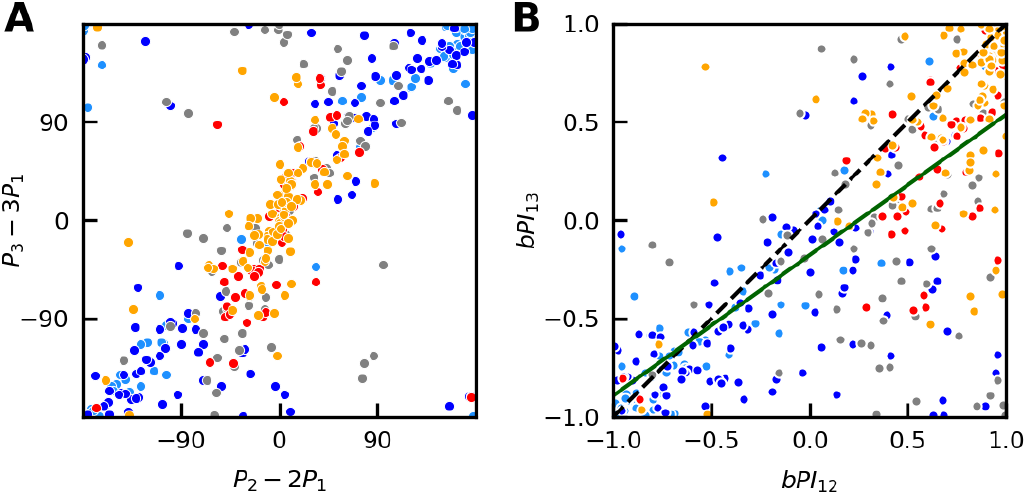
Bikinetic pattern index. **A**: The relationship between the phase of the first and second Fourier harmonic of the tuning curves for type I plaids (abscissa) and that between the phase of the first and third harmonic (ordinate), both carry information about the nature of each cell. For both measures, large values are associated with component cells, and small values with pattern cells. **B**: The *bPI*_12_ and *bPI*_13_ indices are significantly correlated (dark green line, *r* = 0.728, *p* = 5.8 × 10^−65^). Their individual performance can be improved by computing a weighted averaging of them, yielding the *bPI* index.

The cross-validated classification performance of this index, based on our manual classification, is 91.5%, with *d*′ = 2.55. Note that, unlike the original pattern index, it is based exclusively on the direction tuning curve to type I plaids, it does not require measuring responses to 1-D stimuli. It is close to −1 when the tuning curve to type I plaids is bimodal, and it is close to +1 when it is unimodal.

We can now compare this new index to the original pattern index. In Figure 16A we divided our cells in three groups: those with a bimodal tuning curve to 1-D drifting patterns (orange), those for which there is a difference of more than 15 between the preferred direction for 1-D and 2-D drifting patterns (pink), and the remaining cells (gray). This last group represents cells for which the two measures should both be appropriate to characterize their component/pattern character. Indeed, for these cells the two variables are strongly correlated (gray line, *r* = 0.74, *p* = 10^−34^), with little bias (the regression line crosses *bPI* = 0 around *PI* = 0) and good coverage (the *bPI* covers the *PI* range between −6 and 6). For bimodal cells there is however no correlation between the two (orange regression line); the *bPI* correctly classifies them as pattern cells, whereas *PI* mostly classifies them as either component or unclassed. Finally, for the group with unreliable responses to 1-D stimuli the two measures are correlated (pink regression line, *r* = 0.36, *p* = 2.85 × 10^−4^) but there is a clear bias, with many of these cells being classified by the *PI* as component cells but by the *bPI* as pattern cells.

**Figure 16.**
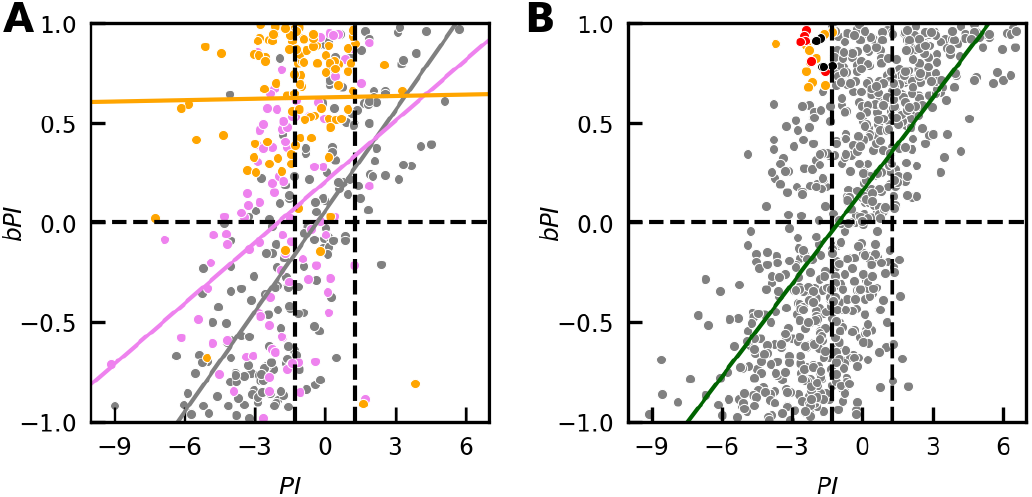
Bikinetic pattern index. **A**: The *bPI* and the original *PI* are correlated. However, this correlation does not hold for cells that were classified as pattern-slow (orange). It is also weaker in cells for which there is a sizable difference in preferred direction to 1-D and 2-D noise patterns (pink), relative to all other cells (gray). **B**: Relationship between *bPI* and *PI* in MT single units recorded using sinusoidal gratings (Wang and Movshon, 2016). With speed stimulus optimized for each cell the correlation between the two is strong. Also in this case a fraction of the cells is classified as component by the *PI* and as pattern by the *bPI*. Colored and black dots referred to cells whose tuning curves are shown in Figure 17.

To further validate this measure, we applied it to a large set of recordings with single electrodes (792 neurons), using sinusoidal gratings and type I plaids as stimuli (Wang and Movshon, 2016). In Figure 16B we plot the *bPI* against the pattern index for these recordings. Also in this case there is a strong correlation between the two measures (dark green line, *r* = 0.635, *p* = 9.53 × 10^−91^), although the regression line is shifted to the left compared to the gray regression line in Figure 16A. The reason for this shift is the presence of a sizable number of cells in the upper-left corner of the plot (i.e., cells classified as component by the *PI* but as pattern by the *bPI*). In Figure 16A cells in that region of the plot had (with rare exceptions) either bimodal responses to the 1-D noise (orange) or unreliable responses to 1-D stimuli (pink). In Figure 17 we plot the tuning curves for the 24 cells from this group with the highest *bPI*. Responses to the sine grating are shown in black, and those to the type I sinusoidal plaid in purple; for some cells responses to 2-D noise stimuli were also available (gray). Obviously, in all cells the response to the type I plaid is mostly unimodal (hence the large *bPI*), and thus more similar to what expected from a pattern cell than from a component cell. In 12 cells (orange numerals in the top left corner of each panel, indicated in orange in Figure 16B) the tuning curve to the sine grating is clearly bimodal, indicating that the speed was probably low relative to the preferred speed of the cell. In 7 cells (red numerals, indicated in red in Figure 16B) the preferred direction to sine gratings and type I plaids coincides, but there is no response to the second grating in the plaid (the type I tuning curves are unimodal). In the three cells (13, 17 and 19) in which the response to random dots stimuli was also recorded, the preferred direction for those stimuli is clearly different from that for sine gratings. If that is indeed the preferred direction for the cell, the response to a type I plaid is exactly what would be expected from a pattern cell. Finally, in the last five cells (black numerals, indicated in black in Figure 16B) the preferred direction for type I and sine gratings is quite different, but there is no response to the first grating in the type I plaid. Again, in two cells (20 and 24) for which random dots responses were recorded, the preferred direction to those stimuli does not match that to sine gratings, but is exactly what would be expected from a pattern cell with the given response to a type I plaid. Thus, even with single units recordings in some cells we observe the same type of responses to 1-D stimuli (random line stimuli in our case, sine gratings for these data) that impair the ability of the pattern index to correctly characterize the cell. In contrast, our new measure is robust to these problems, and it is thus preferable regardless of the stimulus used.

**Figure 17.**
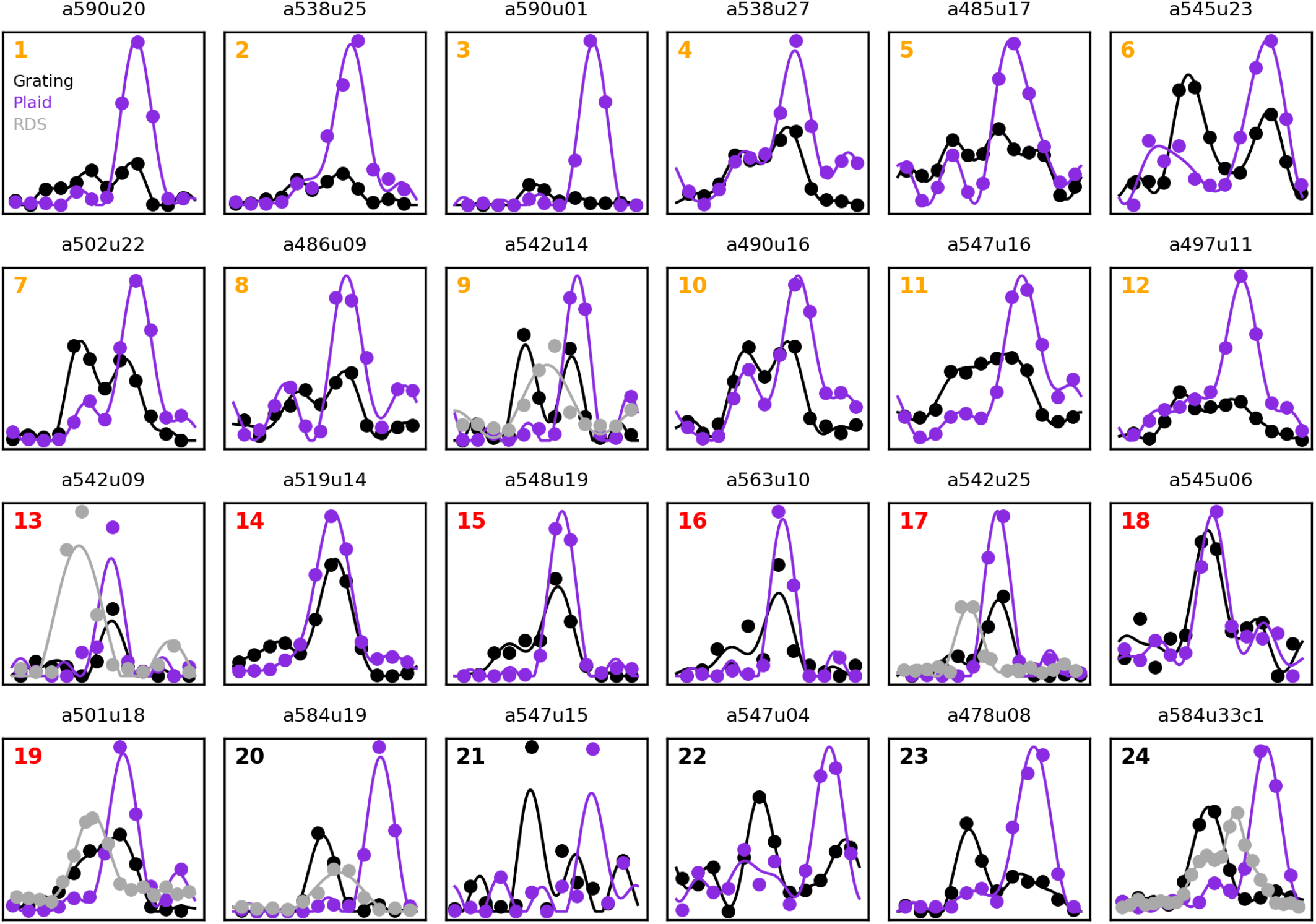
Directional tuning curves to sinusoidal gratings (black), type I sinusoidal plaids (purple), and 2-D noise (gray; only present in few cells, and in some cases with finer directional sampling) for 24 MT cells (Wang and Movshon, 2016). The color of the numeral in each panel is also used to identify the cells in Figure 16B.

### Unikinetic Pattern Index (*uPI*)

With unikinetic plaids, comparing the tuning curves for UP45 and UP-45 plaids is likely the best approach to tell apart component cells from pattern cells. With our approach that means looking at the phase difference in individual harmonics across the two plaids. If we now go back to Figure 14 and look at the tuning curves to unikinetic plaids (center and right columns), we note that, as the peak of the tuning curve (blue) moves away from 0 as we go from component to pattern cells, *P*_1_ (green) barely budges. What could be thought as being the most important harmonic thus seems to be of limited value. Similarly, *P*_3_ (pink) goes through a 180° shift from component to pattern cells, but this is true for both UP45 and UP-45 plaids. The most promising component is then *H*_2_ (red). In component cells, the difference between *P*_2_ in UP45 and UP-45 plaids is rather small, around zero; however, in pattern cells it is large, around 180°. This measure might thus be used to discriminate component from pattern cells. We had also mentioned above that some of the issues with correctly classifying cells with unikinetic plaids stem from their response to the static component. One simple way of attenuating the impact of such responses is to compute, for each cell, the tuning curve of a putative opponent cell (i.e., a cell with a tuning curve that is rotated by 180°), and subtract the latter from the former (clipping negative values to zero). We can then extract the difference between *P*_2_ in UP45 and UP-45 plaids from these opponent tuning curves.

In Figure 18A, we plot, for all cells (color-coded according to our manual classification, as in Figure 15), these two phase differences (single cells on the abscissa, opponent pair on the ordinate). As expected from Figure 14, in component cells (blue and cyan) both of these have values around zero (although in the opponent pairs they are biased toward positive values, whereas in pattern cells - red and orange - both have values around 180°). Like we did above, we can define an index based on each of these two measures:

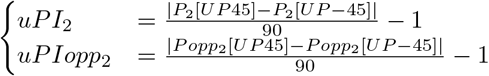

*uPI* stands for unikinetic pattern index. Each index varies between −1 (putative component cell) and 1 (putative pattern cell). In Figure 18B we plot these two indices. They are highly correlated (*r* = 0.729, *p* = 2.9 × 10^−65^), but provide some amount of complementary information, and thus can be averaged into a single index:

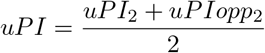

**Figure 18.**
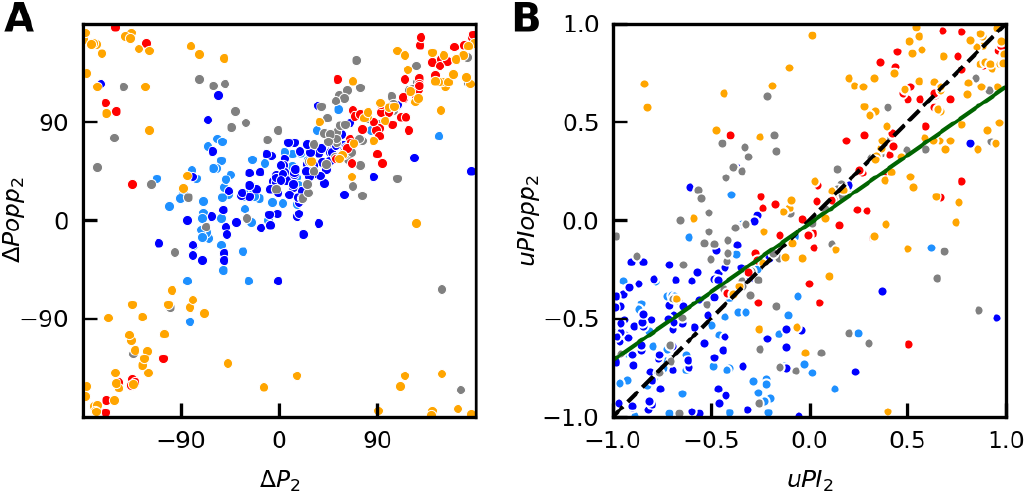
Unikinetic pattern index. **A**: The difference between the phase of the second Fourier harmonic of the tuning curves for UP45 and UP-45 unikinetic plaids (*P*_2_) and that between the same difference for a putative opponent pair (*Popp*_2_) both carry information about the nature of each cell. For both measures, small values are associated with component cells, and large values with pattern cells. **B**: The *uPIopp*_2_ and *uPI*_2_ indices are significantly correlated (dark green line, *r* = 0.729, *p* = 2.9 × 10^−65^). Their individual performance can be improved by averaging them, yielding the *uPI* index.

The cross-validated classification performance of this index, based on our manual classification, is 90.5%, with *d*′ = 2.76. Like the unikinetic rotation it compares responses to UP45 and UP-45 plaids, but instead of directly cross-correlating the tuning curves, it uses only the most sensitive Fourier component, and is augmented using the tuning curve of a putative opponent cell.

### Global Pattern Index (*gPI*)

The *bPI* and *uPI* indices, i.e., our estimates of the “patterness” of a cell based on type I and unikinetic plaids, respectively, thus perform remarkably similarly. They are quite strongly correlated (*r* = 0.61, *p* = 4.2 × 10^−41^, dark green line Figure 19).

**Figure 19.**
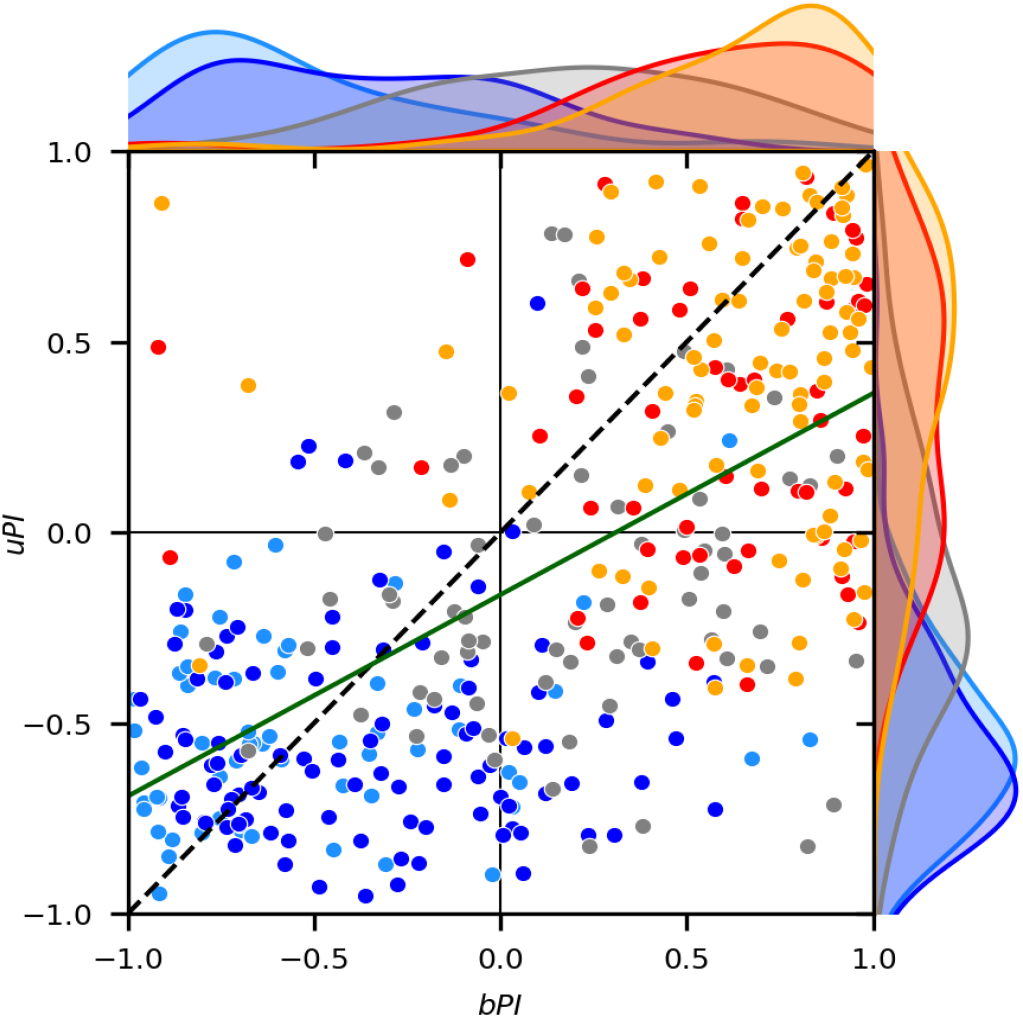
The *bPI* and *uPI* both carry information about the nature of each cell, based on responses to type I and unikinetic plaids, respectively. They are significantly correlated (dark green line, *r* = 0.61, *p* = 4.2 × 10^−41^). Their individual performance can be improved by averaging them, yielding the *gPI* index.

Note that most cells are either classified as component by both indices (third quadrant, *bPI* < 0 and *uPI* < 0), or as pattern by both indices (first quadrant, *bPI* > 0 and *uPI* > 0). There are very few cells that would be classified as pattern based on the *uPI* and as component cells based on the *bPI* (second quadrant), but there is a sizable number of cells that would be classified as component based on the *uPI* and as pattern cells based on the *bPI* (fourth quadrant). An even better alignment with our manual classification can be obtained by simply averaging them into a global pattern index (*gPI*):

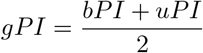

with a cross-validated classification performance equal to 94.0%, with *d*′ = 3.55.

In Figure 20 we plot the distribution of these last three indices for the various classes of cells. We see that the *bPI* does a better job with pattern cells (almost all are associated with a positive value for *bPI*) than with component cells (most have a negative value, but some are positive), and mixed cells have a positive bias. Conversely, *uPI* does a better job with component cells (almost all have a negative value) than with pattern cells (most have a positive value, but some are negative), and mixed cells have a negative bias. The *gPI* is the best of both worlds, assigning negative values to almost all component cells, positive values to almost all pattern cells, and it has zero bias for mixed cells. Note that in all cases the indices do not discriminate between the component and component-fast groups, nor between the pattern and pattern-slow groups, indicating that, as desired, they are insensitive to the speed of the stimulus relative to the preferred speed of the cells. This was not true for the original measures, especially for the pattern index (Figure 2).

**Figure 20.**
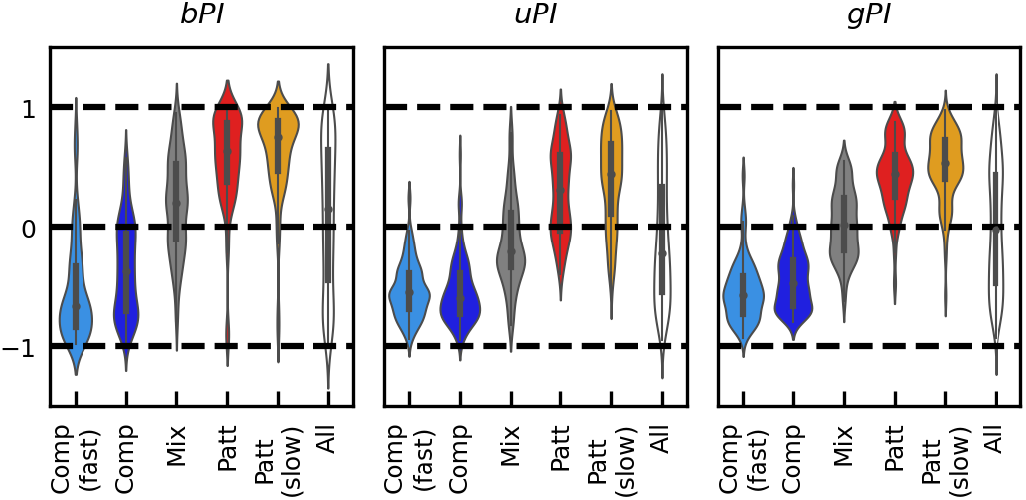
Distribution across our population of neurons of values for three of our proposed indices: bPI (based on the type I plaid tuning curve), uPI (based on the unikinetic plaids tuning curves), and gPI, the average of the two.

It is important to note that the indices that we have developed were not in any way fitted to our manually classified data. Like the pattern index and unikinetic rotation, they are derived from principles of what component and pattern cells should look like, and were only inspired by and validated with our manual classification. Similarly, the combined indices were not fitted to the data to maximize classification performance, using instead fixed combination rules.

### Other single-unit measures

It has been previously reported that, with sinusoidal gratings, the direction bandwidth increases from component to pattern cells. Here we confirm that this holds also with 1-D noise patterns (mean FWHH bandwidth 82° for component cells, 107° for pattern cells, Kolmogorov-Smirnov test, *p* = 0.0068). In contrast, we found that the direction bandwidth for 2-D noise pattern decreases significantly (Kolmogorov-Smirnov test, *p* = 0.0014) from component (120°) to pattern (99.7°) cells (Figure 21). Pattern cells have similar bandwidths for both stimuli (mean log ratio= –0.05), and it is the component cells that have considerably broader 2-D noise than 1-D noise tuning bandwidth (mean log ratio= 0.556, 47% broader). The difference is highly significant (Kolmogorov-Smirnov test, *p* = 1.79 × 10^8^). Here we have considered only the component and pattern groups proper, because measures for the component-fast and pattern-slow groups would be hard to interpret given their bimodal tuning curves.

**Figure 21.**
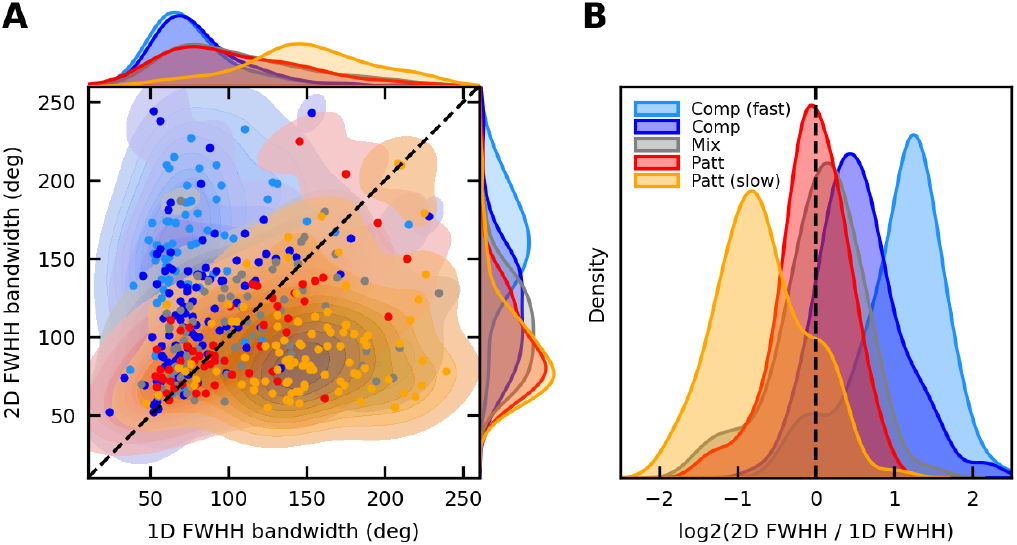
Direction bandwidth for drifting 1-D and 2-D noise patterns for individual cells, color-coded according to our manual classification. Note that the 1-D bandwidth is overestimated in the pattern-slow group, and the 2-D bandwidth is overestimated in the component-fast group. **A**: Scatter plot of the two measures, together with group marginal distributions. **B**: Distribution of the base-2 logarithm of their ratio.

We also found (Figure 22) that 1-D noise patterns are slightly more effective than 2-D noise patterns at driving component cells (on average by 25%), whereas 2-D noise is considerably more effective than 1-D noise at driving pattern cells (on average by 107%). The difference is highly significant (Kolmogorov-Smirnov test, *p* = 9.1*e*14). It thus appears that 1-D noise is a poor stimulus for pattern cells. Since responses to 2-D noise are similar across types, orientation broadband stimuli should be the preferred stimulus for studying MT neurons (when possible).

**Figure 22.**
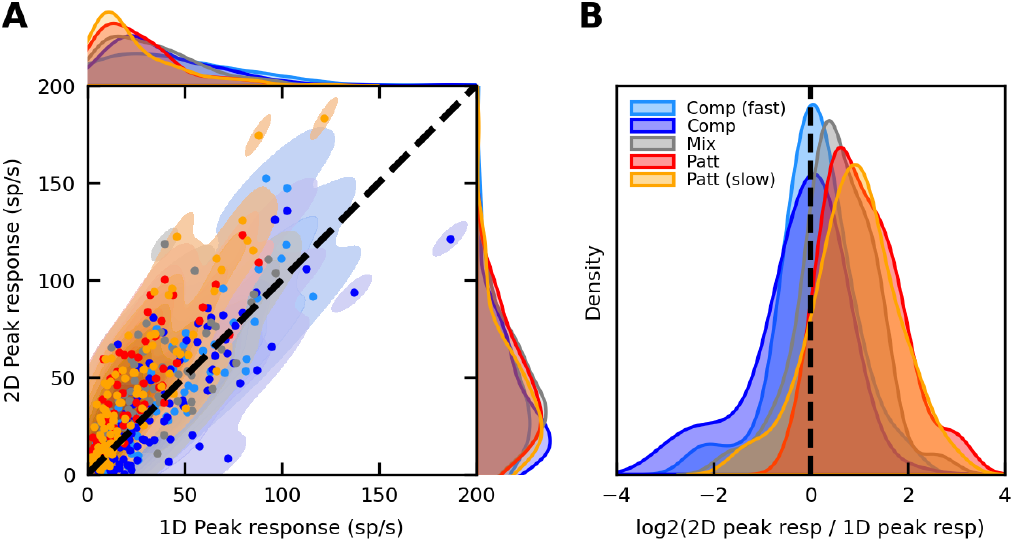
Maximum firing rate for drifting 1-D and 2-D noise patterns for individual cells, color-coded according to our manual classification. **A**: Scatter plot of the two measures, together with group marginal distributions. **B**: Distribution of the base-2 logarithm of their ratio. Pattern cells respond more strongly to 2-D patterns.

### Simultaneous recordings

Because we recorded from multiple units simultaneously, we can determine how trial-by-trial fluctuations vary across pairs of neurons. 383 out of the 386 units presented above have been recorded simultaneously in groups of 2 to 19, across 60 recording sessions. In Figure 23A we plot the distribution of counts of simultaneously recorded units for each session. In Figure 23B we plot the distribution of the distance between the electrode contacts on which each unit in a pair (1454 pairs in all) were recorded. We used as unit of distance the spacing of the contacts, 50 *μm*, and computed the Euclidean distance between the electrodes, which are arranged in two columns of 12 rows each. Zero distance signifies that both neurons in the pair were recorded on the same contact and isolated by spike sorting based on spike shape, a relatively rare occurrence in our dataset. The fact that most pairs were close is a consequence of our strategy for placing the electrode: we positioned it so that easily identifiable single units were not at the end of the arrays, where small drifts could result in the neuron being no longer visible. Most units are then found on the 10 contacts that are less than 5 distance units (0.25 *mm*) away.

**Figure 23.**
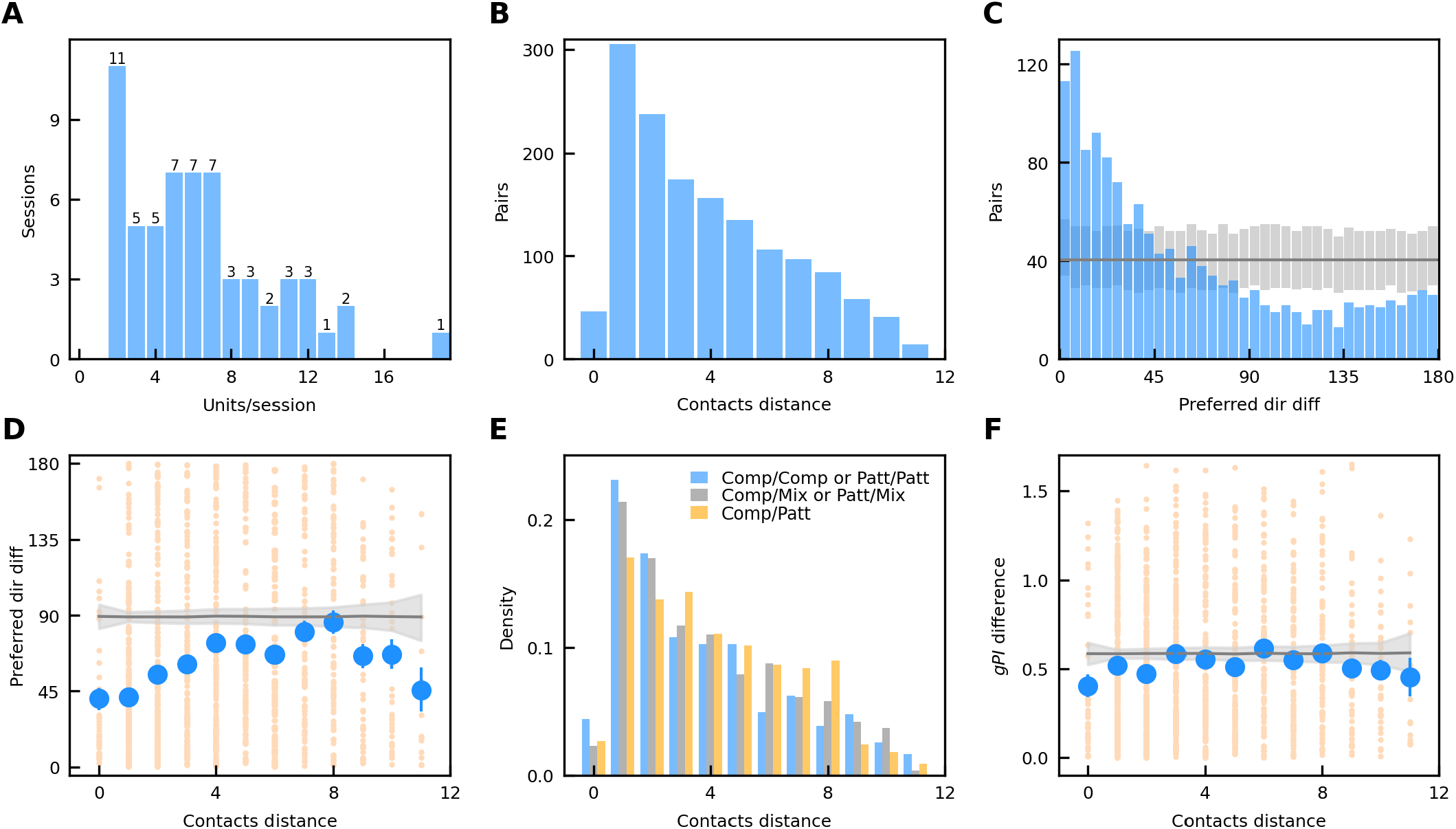
Simultaneous recordings. **A**: Number of units simultaneously recorded in each session (range: 2-19). **B**: Distance between contacts on which the units in each pair were recorded. **C**: Difference in preferred direction between units in a pair (bootstrapped null distribution in gray). **D**: Relationship between contact distance and preferred direction difference. Individual data points in light orange, means for each contact distance (with SEM bars) in blue. The expected mean (with 68% confidence interval shaded in gray) for shuffled data is also shown. **E**: Difference in functional cell type between units in a pair as a function of contacts distance. **F**: Absolute difference in *gPI* between units in a pair as a function of contacts distance. Same conventions as in panel D.

In Figure 23C we plot the distribution of the difference between the preferred directions of the two units in each pair. In our dataset we found a clear preponderance of pairs with cells tuned to similar directions of motion, a distribution that is highly different from that expected by a random sampling of the preferred directions of our 383 units (plotted in gray, 95% confidence interval based on 1000 bootstrapped histograms). Conversely, pairs with a difference larger than 90° are significantly underrepresented. Since most of our pairs are close together (Figure 23B) this reflects the columnar structure for preferred direction in MT (Albright, Desimone, and Gross, 1984). To assess the significance of this deviation from random sampling we have also computed the area under the curve of the cumulative histogram (173,015 for our sample, 95% CI for random sampling: 128,000-135,698), the count of pairs with a difference smaller than 10° (238 in our sample, 95% CI of random sampling: 69-103), and the count of pairs with a difference smaller than 20° (415 in our sample, 95% CI of random sampling: 145-192).

In Figure 23D we show how absolute differences in preferred direction vary with physical distance between the cells in a pair. We found that with small and large distances the mean difference in preferred direction (blue dots; 68% confidence intervals are also plotted, but are often too small to be visible) is considerably smaller than expected by chance (black horizontal line and shaded SEM area, computed by bootstrap analysis of shuffled data): Nearby and far apart cells tend to prefer similar directions of motion. For intermediate distances, between 4 and 8 contact units (0.2 – 0.4 *mm*), there is no more systematic relationship between distance and preferred direction than would be expected by chance. Given that by 11 contact units the mean is similar to that observed at 0 distance (i.e., pairs recorded on the same contact) we infer that, with the angle of our penetrations in MT, the spatial scale for direction is approximately 0.6 *mm*.

To infer whether component and pattern cells are clustered we divided our 386 cells into three groups: Component (n=151) if *gPI* < 0.25, Mixed (n=95) if 0.25 < *gPI* < 0.25, and Pattern (n=140) if *gPI* > 0.25. We then looked at the 1454 pairs of these cells that were recorded simultaneously, and how the contacts distance relates to the ”type distance” (which we defined as 0 if two cells belong to the same group, 1 if one is Mixed and the other Pattern or Component, and 2 if one is Pattern and the other Component). We found (Figure 23E) that the effect of type distance is rather small: For distance 0, the mean contact distance is 3.81 (median 3.0); for distance 1, the mean is 4.03 (median 3.16); for distance 2, the mean is 4.22 (median 4.0). Nevertheless, the effect is significant (Kruskal-Wallis H test, *p* = 0.011), as is the difference between the distribution of contact distances for cells that belong to the same group (type distance 0) as opposed to the opposite group (type distance 2), Mann-Whitney test *p* = 0.0032. In Figure 23F, similarly to what we did in Figure 23D for the preferred direction, we show how absolute differences in *gPI* vary with physical distance between the units in a pair. The difference in *gPI* is intended as a proxy for the functional difference between the cells, expressed in a continuous as opposed to categorical manner. We also show (black horizontal line and shaded gray area) the *gPI* difference that would be expected by randomly drawing cells without regards to their distance. We found that at all distances the mean *gPI* difference (blue dots) lies very close to the chance line, again showing that we found no convincing evidence of clustering by functional cell type.

As previously reported (Zohary, Shadlen, and Newsome, 1994; Cohen and Newsome, 2008) we found (Figure 24) that both signal (panel A, slope: −0.88) and noise (panel B, slope: −0.38) correlations decrease significantly (p < 10^−40^ in both cases) as the difference between the preferred direction of the two neurons increases. This is to be expected, since neurons that signal different directions of motion are bound to share few inputs. For pairs with a difference in preferred direction up to 20° (415), the mean noise correlation is 0.182 (median 0.162, standard deviation 0.157). The fraction of pairs that have noise correlations that are significantly higher than would be expected by chance (95% confidence interval on trial shuffled data) also decreases with the difference in preferred direction, whereas the fraction with noise correlations lower than chance increases (Figure 24C).

**Figure 24.**
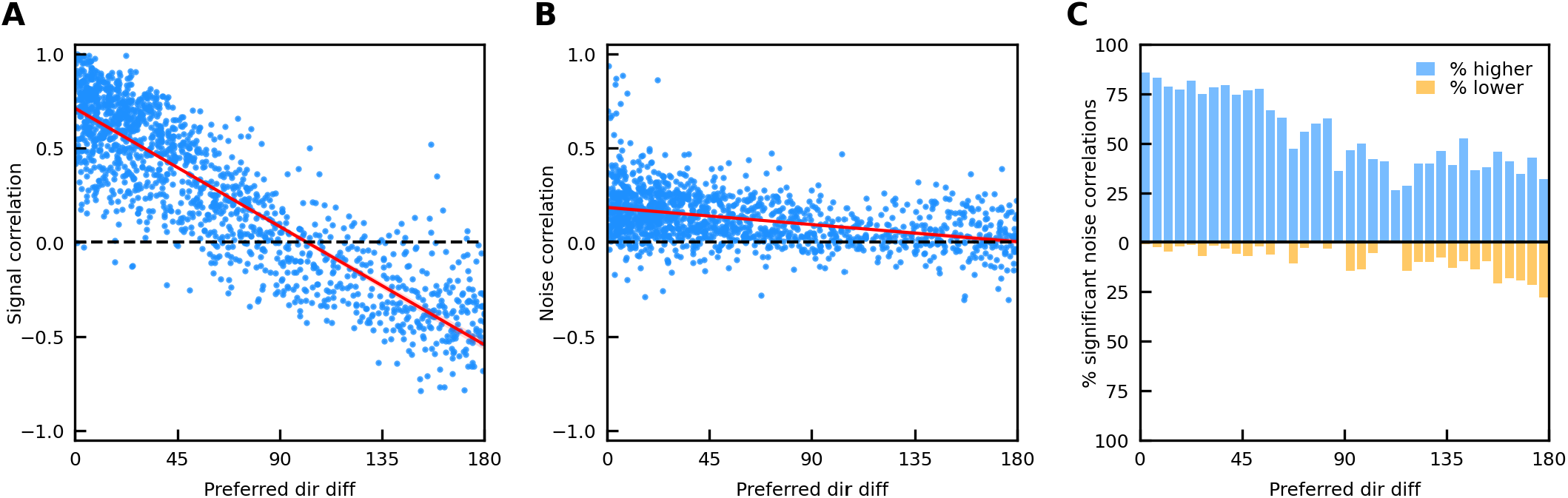
Signal and noise correlations. **A**: Signal correlations decrease with difference in preferred direction, going from strongly positive around 0° to strongly negative around 180°. **B**: Noise correlations also decrease with difference in preferred direction, from positive around 0° to zero around 180°. **C**: When the difference in preferred direction is small, most pairs have a higher noise correlation than expected by chance (blue bars). This percentage drops as the difference increases, whereas that of correlations lower than expected by chance (orange bars) increases.

Finally, we attempted to quantify whether noise correlations are different within subsets of neurons, such as pattern and component cells. Because the difference between the preferred directions of the two cells in a pair has such a strong impact on noise correlations (Figure 24B), we restricted our analysis to pairs in which the difference in preferred directions was smaller than 30° (n=543), so that differences in this parameter across groups would not contaminate the results.

In Figure 25A we show how mean noise correlations vary when pairs are sorted based on our manual cell classification. We also tested other grouping schemes, and the results are always slightly different. In Figure 25B, we show (shading) the local mean noise correlation (computed using the Lowess 2-D smoothing algorithm) as a function of the value of *gPI* for the two neurons in each pair (blue dots)^4^. There is little structure to the correlations, but points close to the identity line, where pairs that have similar values of *gPI* (and thus most likely similar tuning curves to type I and unikinetic plaids) lie, tend to have higher noise correlations than points that lie far from it.

**Figure 25.**
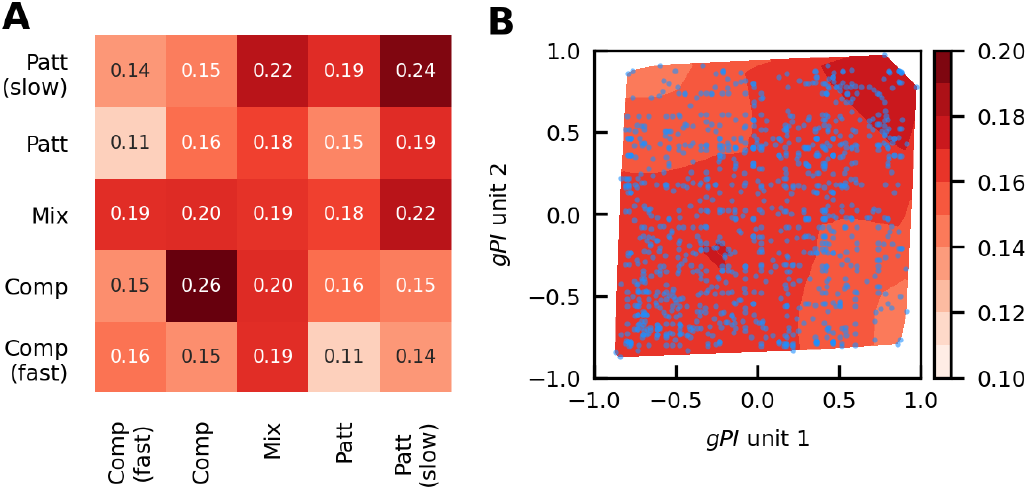
Noise correlations and functional types. Only pairs with a difference in preferred direction smaller than 30° were used in these analyses. **A**: Noise correlation matrix for our manual classification. **B**: Noise correlation averages (shading) as a function of the value of *gPI* for the two neurons in a pair (blue dots).

In summary, while there is statistical evidence that some clustering exists, it is much weaker than that based on preferred direction. We suspect that, at least in a passive fixation task, where noise correlations are probably determined by common bottom-up inputs, what matters most is the similarity of tuning curves, more than any functional grouping.

## Discussion

We described neurophysiological signals recorded from a large number of neurons from area MT of passively fixating monkeys. The stimuli we used, based on random line patterns, are quite different from those used in previous studies, but have several advantages, especially for multi-contact array recordings. Their ability to effectively stimulate a large fraction of cells cannot however overcome the long-known issues (Kawakami and Okamoto, 1996; Simoncelli and Heeger, 1998) associated with presenting stimuli that are not optimized for the preferred speed of a cell.

We showed that stimulating a cell at a suboptimal speed severely limits the ability of previously developed indices to functionally classify MT neurons along the component-pattern axis. In fact, our analyses reveal that even single-unit recordings are not immune to this problem (Figure 17), probably because time constraints inevitably result in a process of stimulus speed optimization that is rather coarse.

To overcome the limits of the existing indices, we introduced two new indices, one (*bPI*) based on the direction tuning curve to type I plaids, and the other (*uPI*) based on the direction tuning curves to two sets of unikinetic plaids. The indices are computed from (some of) the coefficients of the discrete Fourier transform of the tuning curves. The formulas given for the indices are valid for direction tuning curves sampled every 30°, which is a common practice in neurophysiological recordings in MT; different sampling schemes would naturally require the formulas to be adapted.

The major advantage of these new indices is that they perform equally well for optimal and suboptimal stimulus speeds. This is possible because the new indices rely exclusively on the response of the cell to plaids, without attempting to model this response as a function of the responses to the individual components in the plaid. By properly selecting individual Fourier components we were also able to focus on those that are less affected by noise and by the large variability in tuning curve patterns observed across cells. A second advantage of our new indices is that they vary in a fixed range [-1, 1], similarly to indices commonly used to describe a cell sensitivity to other stimulus properties (e.g., orientation, direction of motion, binocular disparity).

### Are there component and pattern subgroups in MT?

Whenever indices are developed to characterize the properties of a cell, the natural question to ask is whether the cells form a continuum or there are distinct groups. What is most striking about the large number of direction tuning curves we have illustrated (Figures 3, 4, 6, 7, 9, 10) is how diverse they are, even for cells that (putatively) belong to the same functional class. Such variability, leading to tuning curves that look quite different from the prototype component and pattern cells (Figure 1), is probably related to each MT cell receiving only a subset of the V1 inputs required to construct an ideal pattern cell. In the context of the Simoncelli and Heeger (1998) model, this would result in only a partial coverage of the velocity plane.

The requirements necessary to construct an ideal pattern cell from component-like (i.e., V1) inputs appear to be more frequently violated for unikinetic than for type I plaids. This is highlighted by the finding that when we plot our index of cell ”patterness” based on unikinetic plaids (*uPI*) versus that based on type I plaids (*bPI*), we find (Figure 19) that while most cells fall in the first (pattern classification based on both stimuli) and third (component classification based on both stimuli) quadrants, many cells fall in the fourth quadrant (pattern-like with type I plaids, component-like with unikinetic plaids), but very few fall in the second quadrant. This replicates previous findings (Wallisch and Movshon, 2019) based on single-unit recordings and sinusoidal plaids (note that to match our sign conventions their Figure 4 must be reflected around the horizontal axis).

We thus suggest that the dichotomy between a continuum and distinct groups is ill posed: it is neither. A more apt metaphor is that of an orchestra. Each player (cell) makes a distinctive contribution, but there are so many different instruments that no single dichotomy provides a valuable classification (woodwind is different from brass, but it is not meaningful to classify each player along a woodwind-brass axis). Having neurons that carry such mixed signals has important computational advantages, enabling flexible readouts (Fusi, Miller, and Rigotti, 2016).

All indices that purport to quantify where a cell lies along a continuum must thus be taken with a grain of salt. As we show in Figures 12 and 13, and as we will discuss at length elsewhere, by listening to a large enough subset of MT neurons, read-out neurons in other areas can hear a distinct melody. To decode pattern and component motion signals in the visual input, downstream areas thus simply need to gather inputs from the appropriate subsets of MT neurons, in spite of individual MT neurons carrying rather muddy signals. The much clearer motion signals observed in area MST (Khawaja, Liu, and Pack, 2013) might thus be the result of simple averaging of MT outputs. Averaging over different subpopulations might allow to extract signals useful for other purposes, such as identifying transparent motion.

## Acknowledgments

This project was supported by the Intramural Research Program of the National Eye Institute, NIH, DHHS.

## Appendix

**Table 1.** Here we report the values of all the indices discussed above for all the sample cells whose tuning curves we have shown in the paper.

| Figure | Cell | $PI$ | $UR$ | $bPI$ | $uPI$ | $gPI$ |
| --- | --- | --- | --- | --- | --- | --- |
| 3 | M2 S7 C5 | -6.14 | 99.10 | 0.58 | 0.18 | 0.38 |
|  | M2 S41 C30 | -5.50 | 57.16 | 0.42 | 0.92 | 0.67 |
|  | M2 S39 C12 | -4.43 | 70.04 | 0.85 | 0.87 | 0.86 |
|  | M1 S166 C8 | -2.97 | 68.46 | 0.25 | 0.59 | 0.42 |
|  | M1 S148 C4 | -2.25 | 50.97 | 0.91 | 0.85 | 0.88 |
| 4 | M1 S138 C19 | -1.60 | 54.53 | 0.48 | 0.58 | 0.53 |
|  | M2 S66 C7 | -1.79 | 85.54 | 0.51 | 0.64 | 0.57 |
|  | M2 S30 C6 | -2.40 | 53.04 | 0.30 | 0.63 | 0.46 |
| 6 | M1 S148 C4 | -2.25 | 50.97 | 0.91 | 0.85 | 0.88 |
|  | M2 S39 C12 | -4.43 | 70.04 | 0.85 | 0.87 | 0.86 |
|  | M2 S38 C2 | -2.80 | 92.51 | 0.93 | 0.58 | 0.75 |
|  | M2 S44 C4 | -3.05 | 112.34 | 0.85 | -0.01 | 0.42 |
| 7 | M1 S188 C2 | -0.51 | 22.20 | 0.95 | -0.23 | 0.36 |
|  | M2 S36 C1 | -2.22 | 44.31 | 0.94 | 0.48 | 0.71 |
|  | M1 S135 C2 | -2.48 | 55.30 | 0.41 | -0.30 | 0.05 |
|  | M1 S183 C6 | -1.39 | 42.35 | 0.53 | -0.06 | 0.24 |
|  | M2 S38 C33 | -7.25 | 47.60 | 0.02 | 0.37 | 0.19 |
|  | M1 S184 C9 | -1.71 | 43.09 | 0.98 | -0.15 | 0.41 |
| 9 | M2 S7 C23 | -6.86 | 26.54 | -0.09 | -0.41 | -0.25 |
|  | M1 S164 C5 | -5.01 | 35.15 | -0.33 | -0.52 | -0.43 |
|  | M2 S33 C14 | -4.04 | 23.71 | -0.09 | -0.53 | -0.31 |
|  | M1 S134 C3 | -3.14 | 34.74 | -0.86 | -0.37 | -0.62 |
|  | M1 S129 C7 | -2.86 | 25.68 | -0.77 | -0.38 | -0.57 |
| 10 | M2 S26 C8 | -5.36 | 50.60 | -0.67 | -0.55 | -0.61 |
|  | M1 S134 C5 | -3.37 | 66.23 | -0.68 | -0.55 | -0.61 |
|  | M1 S105 C21 | -9.13 | 39.82 | -0.71 | -0.38 | -0.55 |
|  | M1 S150 C4 | -2.91 | 51.14 | 0.39 | -0.34 | 0.03 |
|  | M2 S7 C7 | -2.39 | 89.73 | -0.62 | -0.53 | -0.58 |
|  | M2 S7 C3 | -5.54 | 89.35 | -0.75 | -0.22 | -0.49 |

## Footnotes

2 An alternative and just as good estimate of the preferred direction is given by the phase of the first harmonic of the Fourier decomposition of the 1-D and 2-D direction tuning curves.

3 It might be argued that these cells actually belong to the pattern-slow group, as in all cases there is some evidence of a secondary peak in the 1-D noise tuning curve. However, these cells did not meet our criteria for bimodality, either because the secondary peak is too small (first two cells), or because the fitting curve has only a saddle, and no central dip (third cell). It should be noted that if the stimulus was indeed too slow, this would mean that 1-D noise responses in pattern cells can produce direction tuning curves that depart considerably from the bimodal symmetry expected, and shown by the cells in Figure 3. This is a question that can only be answered recording with stimuli of multiple speeds, something we did not do.

4 For each pair there are two dots in this figure, symmetric around the main diagonal.

## References

Albright, TD (1984). Direction and orientation selectivity of neurons in visual area MT of the macaque. J Neurophysiol 52: 1106–30.

Albright, TD, R Desimone, and CG Gross (1984). Columnar organization of directionally selective cells in visual area MT of the macaque. J Neurophysiol 51: 16–31.

Allman, J and J Kaas (1971). A representation of the visual field in the caudal third of the middle temporal gyrus of the owl monkey (aotus trivirgatus). Brain Research 31: 85–105.

Cohen, MR and WT Newsome (2008). Contextdependent changes in functional circuitry in visual area MT. Neuron 60: 162–73.

Cragg, BG (1969). The topography of the afferent projections in the circumstriate visual cortex of the monkey studied by the nauta method. Vision Res 9: 733–747.

De Valois, RL, EW Yund, and N Hepler (1982). The orientation and direction selectivity of cells in macaque visual cortex. Vision Res 22: 531–44.

Dubner, R and SM Zeki (1971). Response properties and receptive fields of cells in an anatomically defined region of the superior temporal sulcus in the monkey. Brain Research 35: 528–532.

Fusi, S, EK Miller, and M Rigotti (2016). Why neurons mix: high dimensionality for higher cognition. Curr Opin Neurobiol 37: 66–74.

Hamilton, DB, DG Albrecht, and WS Geisler (1989). Visual cortical receptive fields in monkey and cat: spatial and temporal phase transfer function. Vision Res 29: 1285–308.

Hawken, MJ, AJ Parker, and JS Lund (1988). Laminar organization and contrast sensitivity of direction-selective cells in the striate cortex of the old world monkey. J Neurosci 8: 3541–8.

Kawakami, S and H Okamoto (1996). A cell model for the detection of local image motion on the magnocellular pathway of the visual cortex. Vision Res 36: 117–147.

Khawaja, FA, LD Liu, and CC Pack (2013). Responses of MST neurons to plaid stimuli. J Neurophysiol 110: 63–74.

Kumano, H and T Uka (2013). Responses to random dot motion reveal prevalence of pattern-motion selectivity in area MT. J Neurosci 33: 15161–70.

Maunsell, JH and DC van Essen (1983a). The connections of the middle temporal visual area (MT) and their relationship to a cortical hierarchy in the macaque monkey. J Neurosci 3: 2563–86.

Maunsell, JH and DC Van Essen (1983b). Functional properties of neurons in middle temporal visual area of the macaque monkey. I. Selectivity for stimulus direction, speed, and orientation. J Neurophysiol 49: 1127–47.

Movshon, JA, EH Adelson, MS Gizzi, and WT Newsome (1985). The analysis of moving visual patterns. In Chagas, C, R Gattass, and C Gross, editors, Pattern recognition mechanisms, pp. 117–151, Rome. Vatican Press.

Movshon, JA and WT Newsome (1996). Visual response properties of striate cortical neurons projecting to area MT in macaque monkeys. J Neurosci 16: 7733–41.

Nishimoto, S and JL Gallant (2011). A three-dimensional spatiotemporal receptive field model explains responses of area MT neurons to naturalistic movies. J Neurosci 31: 14551–64.

Okamoto, H, S Kawakami, H Saito, E Hida, K Odajima, D Tamanoi, and H Ohno (1999). MT neurons in the macaque exhibited two types of bimodal direction tuning as predicted by a model for visual motion detection. Vision Res 39: 3465–79.

Perrone, JA (2004). A visual motion sensor based on the properties of V1 and MT neurons. Vision Res 44: 1733–55.

Prince, SJ, AD Pointon, BG Cumming, and AJ Parker (2002). Quantitative analysis of the responses of V1 neurons to horizontal disparity in dynamic random-dot stereograms. J Neurophysiol 87: 191–208.

Quaia, C, EJ FitzGibbon, LM Optican, and BG Cumming (2019). Motion and binocular disparity processing: Two sides of two different coins. Prog Brain Res 248: 157–166.

Quaia, C, LM Optican, and BG Cumming (2016). A motion-from-form mechanism contributes to extracting pattern motion from plaids. J Neurosci 36: 3903–18.

Rust, NC, V Mante, EP Simoncelli, and JA Movshon (2006). How MT cells analyze the motion of visual patterns. Nat Neurosci 9: 1421–31.

Salzman, CD, CM Murasugi, KH Britten, and WT Newsome (1992). Microstimulation in visual area MT: effects on direction discrimination performance. J Neurosci 12: 2331–55.

Simoncelli, EP and DJ Heeger (1998). A model of neuronal responses in visual area MT. Vision Res 38: 743–61.

Smith, MA, NJ Majaj, and JA Movshon (2005). Dynamics of motion signaling by neurons in macaque area MT. Nat Neurosci 8: 220–8.

Ungerleider, LG and M Mishkin (1979). The striate projection zone in the superior temporal sulcus of macaca mulatta: location and topographic organization. J Comp Neurol 188: 347–66.

Van Essen, DC, JH Maunsell, and JL Bixby (1981). The middle temporal visual area in the macaque: myeloarchitecture, connections, functional properties and topographic organization. J Comp Neurol 199: 293–326.

Wallisch, P and JA Movshon (2019). Responses of neurons in macaque mt to unikinetic plaids. J Neurophysiol 122: 1937–1945.

Wang, HX and JA Movshon (2016). Properties of pattern and component direction-selective cells in area MT of the macaque. J Neurophysiol 115: 2705–20.

Zohary, E, MN Shadlen, and WT Newsome (1994). Correlated neuronal discharge rate and its implications for psychophysical performance. Nature 370: 140–3.

